# Identifying dysregulated regions in amyotrophic lateral sclerosis through chromatin accessibility outliers

**DOI:** 10.1101/2023.08.25.554881

**Authors:** Muhammed Hasan Celik, Julien Gagneur, Ryan G Lim, Jie Wu, Leslie M. Thompson, Xiaohui Xie

**Affiliations:** Department of Computer Science, University of California Irvine, Irvine, CA, USA; Center for Complex Biological Systems, University of California Irvine, Irvine, CA, USA; Department of Informatics, Technical University of Munich, Garching, Germany; Helmholtz Association - Munich School for Data Science (MUDS), Munich, Germany; Institute of Human Genetics, School of Medicine, Technical University of Munich, Munich, Germany; Institute of Computational Biology, Helmholtz Center Munich, Neuherberg, Germany; Institute for Memory Impairments and Neurological Disorders, University of California Irvine, Irvine, CA 92697, USA; Department of Biological Chemistry, University of California Irvine, Irvine, CA, USA; UCI MIND, University of California Irvine, Irvine, CA, USA; Department of Psychiatry and Human Behavior and Sue and Bill Gross Stem Cell Center, University of California Irvine, Irvine, CA, USA; Department of Neurobiology and Behavior, University of California Irvine, Irvine, CA, USA

## Abstract

The high heritability of ALS and similar rare diseases contrasts with their low molecular diagnosis rate post-genetic testing, pointing to potential undiscovered genetic factors. Chromatin accessibility assays quantify the activity of functional elements genome-wide, offering invaluable insights into dysregulated regions. In this research, we introduced EpiOut, a computational toolbox to identify outliers in chromatin accessibility. These outliers represent dysregulated regions where chromatin accessibility uniquely diverges from the population baseline in a single or few samples. Annotation of accessible regions with histone ChIP-seq and Hi-C indicates that outliers are concentrated in functional loci, especially among promoters interacting with active enhancers. Across different omics levels, outliers are robustly replicated, and chromatin accessibility outliers are reliable predictors of gene expression outliers and aberrant protein levels. For example, 59% of gene expression outliers can be linked to aberration in chromatin accessibility. When promoter accessibility does not align with gene expression, our results indicate that molecular aberrations are more likely to be linked to post-transcriptional regulation rather than transcriptional regulation. Our findings demonstrate that the outlier detection paradigm can uncover dysregulated regions in rare diseases. EpiOut is open-sourced and freely available at github.com/uci-cbcl/EpiOut.

## Introduction

Amyotrophic lateral sclerosis (ALS) is a rare neuromuscular degenerative disease affecting 0.6 to 3.8 per 100,000 people with a poor survival prognosis without a cure^1,2^. ALS is a complex disease where a single gene or pathway cannot explain the disease phenotype due to the heterogeneity of genetic causes and over 30 genes associated with ALS^1,3^. Meta-analysis and twin studies estimate the heritability of ALS disease at 61% (with 38%-78% confidence intervals) in sporadic cases (sALS), i.e., patients without a history of the disease in the family^4^. Despite the high heritability of ALS, only 11 to 25% of patients^5–7^ receive a diagnosis after genetic testing. The gap between high heritability and low diagnostic rate implies the existence of many undiscovered ALS-related genes.

There are large-scale sequencing efforts to discover the genetic bases of ALS^8,9^. These studies utilized GWAS, QTL, and differential expression analysis from a large cohort of samples to detect aberrations in ALS patients compared to control samples^8,10–12^. These statistical approaches successfully detected the most common factors (variants, genes, and pathways) associated with disease phenotype, yet detecting rare genetic factors remains challenging due to low statistical power. Outlier detection is a complementary statistical approach that uncovers aberrations specific to one or a few patients. Applying the outlier detection paradigm to transcriptomics data has revealed the dysregulation of many novel splicing and gene expression outliers^13–19^. This approach offers a promising direction for enhancing molecular diagnostic rates of rare disorders, as it effectively captures their heterogeneous genetic architecture.

The outlier detection approach has recently been applied to proteomics^20,21^ and methylation^21^, and robust replication of aberrations across multiple omics data demonstrates the reliability of the outliers for disease diagnostics. Expanding the outlier detection approach to chromatin accessibility could provide further insight into the dysregulation of functional regions and their impact on gene expression in disease. Because transcription factors typically bind to open chromatin regions, defining the activity of promoters and enhancers, which in turn regulate transcription^22^. Thus, aberrations in chromatin accessibility correlate with the dysregulation of gene expression potentially linked to diseases^23–25^. Despite the widespread use of assays such as ATAC-seq and DNase-seq enabling genome-wide investigation of the chromatin accessibility landscape^26^, the outlier detection approach has not been applied to chromatin accessibility data to the best of our knowledge.

Here, we present EpiOut, a software developed for chromatin accessibility outlier detection. Our proposed method takes read alignment files and accessible regions as input, performs ultra-fast read counting per accessible region, detects outliers using a linear autoencoder (LR-AE) with a negative binomial objective function, and annotates outlier regions using ChIP-seq and Hi-C. Optimization of the decoder layer and dispersion parameters requires solving a large number of independent convex problems. We significantly accelerated the LR-AE using TensorFlow^27^ by utilizing a vectorized variation of backtracking line search for the dispersion parameters and L-BFGS for the decoder layer. We applied EpiOut to chromatin accessibility data from ALS samples to identify aberrations in chromatin accessibility that might be linked to the disease. EpiOut pinpoints a small number of sample-specific loci as outliers. Comparison of chromatin accessibility outliers with gene expression outliers and protein aberrations reveals consistent replication across multiple omics levels. This analysis can offer valuable insights into whether aberrations in molecular phenotype are influenced by transcriptional or post-transcriptional regulation. The outlier detection approach identifies known ALS genes and potentially novel disease gene candidates.

## Results

In this study, we explored aberrant chromatin accessibility in ALS using ATAC-seq experiments from the AnswerALS dataset (Methods). This dataset comprises paired genomics, transcriptomics, chromatin accessibility, and proteomics experiments from 253 individuals. With our novel method, EpiOut, we pinpointed regions with abnormal chromatin accessibility and investigated the molecular impact of these outliers by comparing paired experiments across different omics levels. Our results highlight the biological relevance and robustness of the identified accessibility outliers.

### Detection of accessible regions

Detection of aberrant accessibility requires read counts for a set of accessible regions consistent across the individuals. First, we merged ATAC-seq reads of all samples into one meta-sample, then performed joint peak calling with MACS2^28^ using the meta-sample to obtain a consistent set of accessible regions. We detected a total of 858,268 peaks before any filtering. Next, we counted the number of ATAC-seq reads overlapping with each accessible region. Counting reads from a large sample cohort is computationally intensive^15^. Thus, we developed EpiCount, an efficient read counter designed for accessibility data. EpiCount is twice as fast as the state-of-the-art counting tool^29^ (Methods), using ∼50times less memory (Extended Data Fig. 1a,b) and fifteen times faster than the reported runtime of commonly used counting methods^15^. The tidy memory footprint of EpiCount enables the embarrassing parallelization of counting across a large number of samples. Lastly, we filtered accessible regions based on the read counts because many accessible regions often do not replicate across samples. We imposed a replication filter to these peaks, ensuring that the accessible regions were observed in at least 50% of the samples with a minimum of two reads and exhibited high accessibility (100 reads) in at least one sample (Extended Data Fig. 2a). Applying these filters yielded 114,428 accessible regions replicated across samples.

### Outlier detection and benchmark

Using an outlier detection approach, we aim to spotlight the rare aberrations unique to a few or single samples. This was achieved by eliminating major covariation in chromatin accessibility data and examining the remaining variance between samples. We investigated the relationship between the principal components of accessibility data and the disease status of the samples. Fig. 1a illustrates the lack of clustering between samples by disease phenotype along the site of the top two principal components of chromatin accessibility. The top 25 principal components account for approximately 94%of the chromatin accessibility variation between samples. However, none of these top principal components significantly separate ALS samples from controls in this cohort (Extended Data Fig. 3a, b). The observations align with the biology of ALS, given that the most prevalent cause, a hexanucleotide (GGGGCC) repeat expansion in the C9orf72, is present in only about 7% of patients, and other known factors account for merely 1-2% of cases^7^. Thus, focusing on the rare aberrations might reveal dysregulation associated with ALS.

**Fig. 1:**
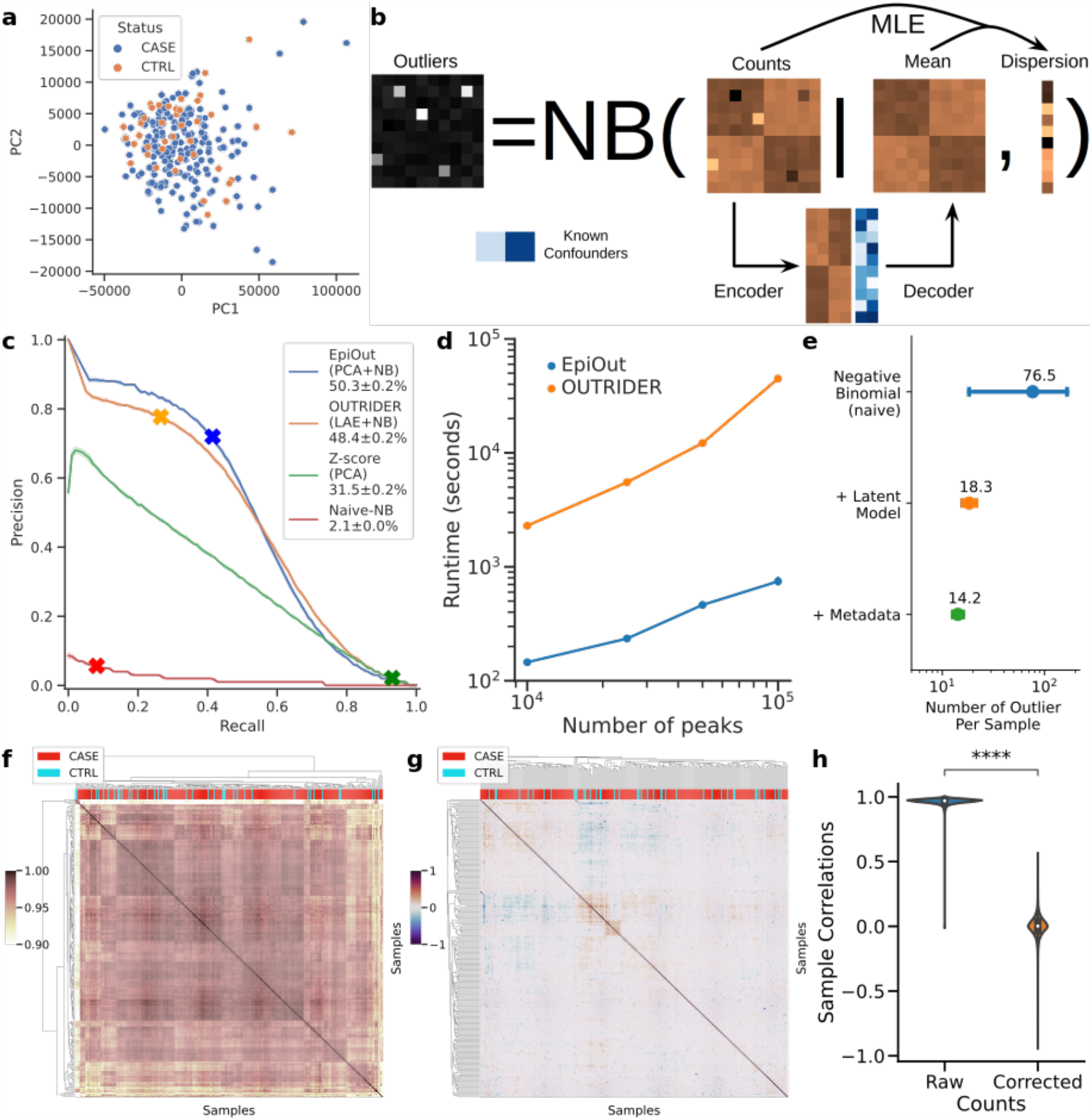
Proposed outlier detection methodology and its benchmark. **(a)** Samples do not form distinct clusters based on their disease phenotype according to the top two principal components of chromatin accessibility. **(b)** The architecture of the proposed methodology for outlier detection (EpiOut). The approach employs the negative binomial test. The mean parameter of the negative binomial distribution is predicted with an LR-AE, which uses latent confounders obtained from data in addition to reported known confounding as features to predict sample-specific expected accessibility. The dispersion parameter is fitted with MLE using the observed and expected counts. **(c)** A precision-recall curve shows the performance of alternative outlier detection methods. Methods were benchmarked based on the classification accuracy of the injected artificial outlier. Specific cutoffs of models are indicated with cross marks (an absolute Z-score of 2 for PCA, a p-value of 0. 05 for naive negative binomial, OUTRIDER, and EpiOut). **(d)** Runtime benchmark of outlier detection methods. **(e)** Contribution of each component (such as latent and known confounding factors) of the model to reduce the number of outliers per sample. **(f)** A Cluster heatmap displaying the cross-correlation of samples based on the raw accessibility counts and **(g)** cross-correlation of samples after correction of counts. **(h)** Distribution of cross-correlation between sample pairs before and after correction of accessibility reads.

To evaluate the performance of the outlier detection methods, we employed an artificial outlier injection procedure previously proposed for detecting aberrant gene expression^30^. To create ground truth, we injected large aberrations, called artificial outliers, to read counts of ALS samples and then benchmarked the performance of tools to classify those artificial outliers on the area under the precision-recall curve (Methods) (Fig. 1c). In naive negative binomial, we estimated the mean of the negative binomial test as a sample mean of read count per ATAC-seq peaks (Methods) and estimated dispersion with maximum likelihood estimation (MLE), then ranked predictions by p-value based on the negative binomial test. The naive negative binomial model performs poorly with area under the precision recall curve (auPRC) of 1.8 ±1%because the expected read counts of the naive negative binomial model are not sample-specific. As an alternative outlier detection method, expected read counts per peak and sample can be estimated by PCA (Methods), and ranking predicted outliers by Z-score based on the expected and observed read counts of the PCA model have the performance of auPRC 9.7 ± 0.9%. Our proposed method estimates expected read counts using an LR-AE (Fig. 1b). Expected read counts are incorporated into a negative binomial test as the mean parameter, and the dispersion is estimated based on the expected and observed counts (Methods). In our implementation of the LR-AE, the weights of the encoder and decoder layers are initialized with the rotation matrix of PCA. Then, the dispersion parameter is initially estimated using expected counts based on initial weights, and weights of the decoder layer are updated to maximize negative binomial likelihood using initial dispersion estimation. After the decoder layer optimization, we recalculate the dispersion estimation and apply the negative binomial test to estimate outliers. Also, it is critical to account for read coverage differences between samples (Extend Data Fig. 2b); thus, read counts are normalized for size factors^31^ (Methods). We chose the optimal bottleneck size of LR-AE with hyperparameter tuning on the validation set (Extended Data Fig. 4). This approach outperforms the previous two methods by achieving an auPRC of 5 0.3 ± 0.2%. An alternative LR-AE-based method, OUTRIDER, outperforms PCA and performs similarly with EpiOut (auPRC of 48 ± 0.2%). Both methods employ an LR-AE with a negative binomial, a more proper distribution to fit counts data than PCA (Methods). Those results show that the estimation of dispersion and gene/sample-specific accessibility expectation followed by the negative binomial test is critical for outlier detection.

Chromatin accessibility is higher dimensional than gene expression because accessibility data may contain hundreds of thousands of accessible genomic regions. In contrast, gene expression data only contains around ten to fifteen thousand expressed protein-coding genes. Thus, the scalability of the outlier detection approach is essential to apply the method to high dimensional chromatin accessibility data. Although OUTRIDER and EpiOut have similar auPRC scores, the autoencoder implementation of OUTRIDER is significantly slower than our proposed method (Fig. 1d). For example, outlier detection with EpiOut (745 ±49 seconds) is 60 times faster than OUTRIDER (44,874 ±200 seconds). Our implementation is faster due to a couple of reasons. Firstly, we do not optimize the encoder layer of the autoencoder because we observed that the initial estimation of encoder weights with PCA is close to optimal, so further training of the encoder is unnecessary. Also, EpiOut only performs one alternating optimization step to fit the decoder layer and estimate dispersion. In contrast, OUTRIDER performs multiple alternative optimization steps to estimate dispersion and train the encoder and decoder layers. Lastly, the optimization of the decoder layer and the estimation of dispersion parameters necessitate solving a multitude of independent convex optimization problems. We efficiently approach these using a vectorized L-BFGS for the decoder layer and a vectorized and bounded backtracking line search for dispersion estimation, all implemented in TensorFlow (Methods).

We further integrated known confounding factors such as batch ID, sex, reported race, and ethnicity into outlier detection (Methods). The contribution of each feature for outlier detection is summarized in Fig. 1e. The naive negative binomial test estimates 76.5 ±614 outlier regions per sample, and the inclusion of latent confounding factors for expected read count estimation significantly reduces the number of outliers to 18.3 ±24. Known confounding factors such as batch ID, sex, race, and ethnicity further reduce the number of an outlier to 14.2 ±13.6 out of 114,428 accessible regions per sample. Thus, our method pinpoints only a handful of aberrant regions per sample.

As another benchmark, we compared cross-correlation between samples. Samples are highly cross-correlated based on the raw read counts (Fig. 1f). In our detection approach, we aim to eliminate the correlation between samples to detect rare aberrations by controlling for latent cofounders. Correcting for latent and known confounding factors and normalization of read counts decorrelate samples and eliminate clusters due to potential batch effects (Fig. 1g). Samples have an average cross-correlation of 96.1%based on the raw read counts. In comparison, there is a 0%average cross-correlation after the proposed correction method (*P* < 0. 001 based on the paired Wilcoxon test, Fig. 1h).

### Functional annotation of accessible regions and outliers

To aid the functional interpretation of chromatin accessibility outliers, we developed EpiAnnot, which annotates accessible regions for histone marks and 3D chromatin interactions based on ChIP-seq and Hi-C experiments. EpiAnnot integrates publicly available ChIP-seq and Hi-C for cell lines/tissues available in Roadmap Epigenomics^32^ and ENCODE^33^ or from custom data sources. We annotated the previously identified accessible regions and outliers using the H3K4me3, H3K27ac, and H3K4me1 histone marks observed in motor neurons. These neurons were derived from the iPSCs of clinically healthy individuals and ALS samples^34^. Based on histone marks, accessible regions were further classified as a promoter if the region has a histone mark of H3K4me3 and is within 1000 base pair (bp) vicinity of the annotated transcript start site or overlaps with 5’ UTR, an active enhancer if the region has both H3K27ac and H3K4me1 mark, and a poised enhancer if only H3K4me1 signal is present while H3K27ac mark is lacking (Fig. 2a)^35^. The two largest categories of accessible regions were active enhancers (n = 39,559), which have both H3K27ac and H3K4me1, followed by poised enhancers (*n* = 18,817) regions, which have only the H3K4me1 mark (Fig. 2b). 14.3%accessible regions (*n* = 16,446) have all three H3K4me3, H3K27ac, and H3K4me1 histone marks. Chromatin accessibility outliers are enriched for histone marks for example 28% of outlier overlap with all three histone marks (Fig. 2c). Consequently, both over-accessibility and under-accessibility outliers are more likely to occur in promoter regions (Fig. 2d, *P* < 0. 001 for both based on Fisher’s exact test). Active enhancers are the largest category of outliers, contain 39% of over-accessibility and 38% of under-accessibility outliers, and not significantly enriched or depleted for outliers (*P* = 0.26 for under-accessibility and *P* = 0.22 for over-accessibility based on Fisher’s exact test). Furthermore, both poised enhancer and unannotated regions strongly depleted for under-accessibility outliers as expected (*P* < 0. 001 for both based on Fisher’s exact test). Significant enrichment of outliers in functional regions indicates the potential utility of accessibility outliers in delineating molecular basis of ALS.

**Fig. 2:**
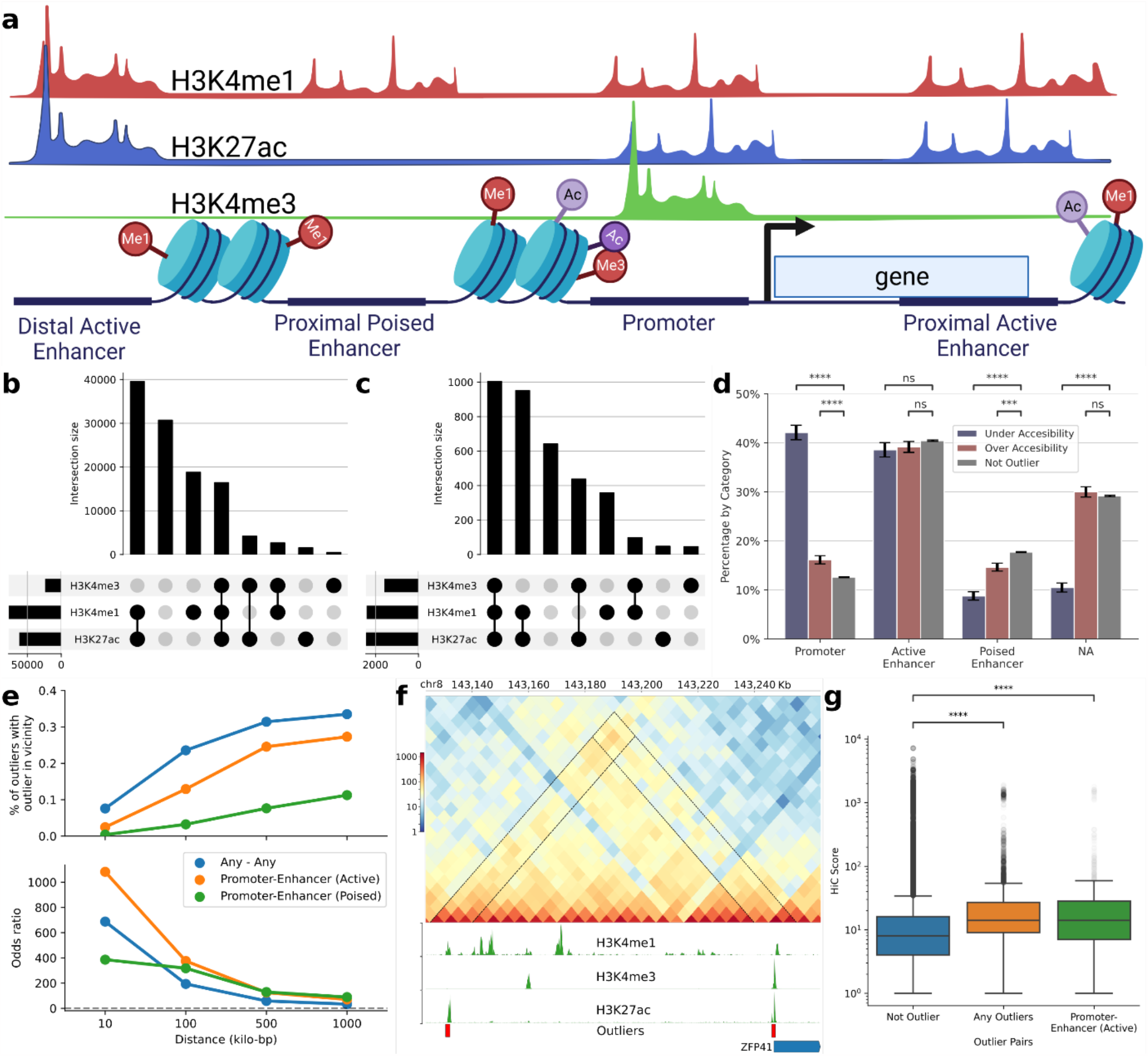
**(a)** Accessible regions were annotated using ChIP-seq marks and gene annotation as promoters, active or poised, and proximal or distal enhancers. **(b)** Overlap between H3K4me3, H3K27ac, and H3K4me1 histone marks and accessible regions **(c)** and outliers **(d)** Breakdown of promoters, active enhancers, poised enhancers, and unannotated regions within the categories of under-accessibility, over-accessibility outliers, and non-outliers. Errors bars indicate standard error and p-values calculated with Fisher’s exact test and corrected for multiple testing with the Benjamini/Hochberg method **(e)** Cumulative odds of observing the second outlier in a region given that there is an outlier in the region (top) and the cumulative percentage of outliers with a second outlier in the vicinity (bottom) based on the distance between regions and the annotation. **(f)** The interaction between the outlier promoter of ZFP41 and ∼1 00 kilo-bp apart outlier distal enhancer is highlighted by the Hi-C track containing contact score between 5 kilo-bp long genomic bins. Coverage tracks for H3K4me3, H3K27ac, and H3K4me1 histone marks are colored green. Red boxes indicate the outlier status of the accessible regions. **(g)** the Hi-C contact scores distribution of non-outlier, outlier, and promoter-enhancer pairs. P-values were calculated with the Mann–Whitney U test and corrected with the Bonferroni correction for multiple testing.

Another interesting observation is that outliers tended to occur in the vicinity of each other (Fig. 2e). Specifically, 8%of outliers have a second outlier in the 10 kilo-bp vicinity with an odds ratio of 688 (*P* < 0. 001 based on the Fisher’s exact test), and 34%of outliers have a second outlier in 1 million bp with the odds ratio of 33 (*P* < 0. 001 based on the Fisher’s exact test). We repeated enrichment analysis between promoter outliers and active or poised enhancer outliers and again observed significant enrichment. 13%of promoter outliers have at least one active enhancer outlier in 100 kilo-bp vicinity (odds ratio = 375, *P* < 0. 001 based on the Fisher’s exact test), and 24%of promoter outliers have an active enhancer outlier in 1 million bp vicinity (odds ratio = 7 0, *P* < 0. 001 based on the Fisher’s exact test). The co-occurrence of the outliers indicates the potential interaction between them. Hi-C experiments from motor neurons provide further evidence for the potential interaction between outliers. For example, the relatively high Hi-C contact score between the outlier promoter of the *ZFP41* gene and a distal enhancer outlier located ∼110 kilo-bp upstream of the gene suggests a potential interaction between outliers (Fig. 2f). We calculated the Hi-C contact score between pairs of accessible regions and categorized regions by outlier status. We observed that outlier pairs (*P* < 0. 001 based on the Mann–Whitney U test), including promoter-active enhancer pairs (*P* < 0. 001 based on the Mann–Whitney U test), have higher interaction scores compared to a baseline where at least one of the regions in the pair is not an outlier (Fig. 2g). Hi-C contact scores are distance-dependent and decay according to power-law with increasing genomic distance. Thus, a higher interaction score could possibly be confounded by the co-occurrence of outliers in the vicinity of each other. To avoid this potential bias, we fit power regression on Hi-C contact scores using the outlier status as a feature and distance as a control variable (Methods). Outlier pairs have higher interaction scores (*P* < 0. 001 based on the t-test) even after controlling for distance with power regression (Extended Data Fig. 5a). Moreover, even when we restricted our analysis to region pairs at least 100,000 bp apart, the Hi-C contact scores of outlier pairs still surpassed the baseline of non-outlier pairs (Extended Data Fig. 5b). Overall, the co-occurrence of outliers in the vicinity of each other and higher Hi-C contact scores between outlier pairs indicate a potential interaction between outliers.

### Chromatin accessibility outliers influence gene expression and protein levels

Aberration in accessibility can impact downstream molecular phenotypes such as gene expression and protein levels. As an example, we found that the ALS case with an under-accessibility outlier in the promoter of *LCMT1* (Fig. 3a-c) also showed decreased mRNA (Fig. 3d) and protein levels (Fig. 3e).

**Fig. 3:**
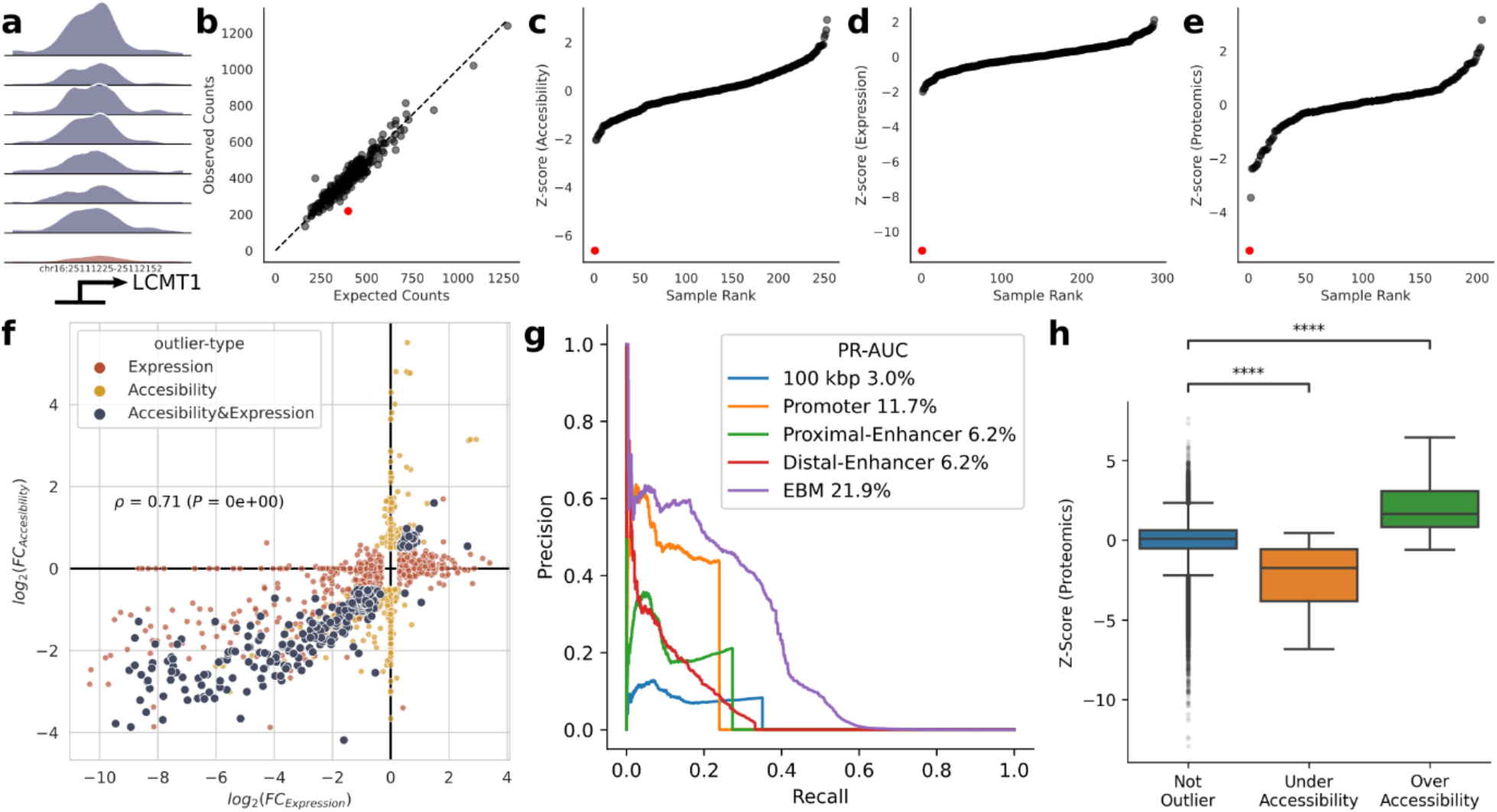
Prediction of gene expression outliers and aberrant protein levels from chromatin accessibility. **(a)** The outlier promoter of the *LCMT1* gene (in red) has a much lower ATAC-seq read coverage than promoters of non-outlier samples. **(b)** Expected and observed accessibility in the promoter of LCMT1 gene across samples. The outlier sample is indicated with a red dot in the figure panels. **(c)** Z-score distribution of promoter accessibility, **(d)** gene expression, **(e)** and protein levels of *LCMT1* across samples. **(f)** Correlation between the absolute log fold change of accessibility and expression outliers. **(g)** The precision-recall curve compares the performance of a range of predictors to estimate gene expression outliers. Those predictors are the absolute log fold change of promoter outliers (orange), the absolute log fold change of proximal enhancers (green), the maximum absolute fold change of any outlier in 100 kilo-bp vicinity (blue), the absolute log fold change of distal enhancers weighted by ABC score (red), explainable boosting machine (EBM) trained with promoter, proximal-enhancer, and distal-enhancers features (purple). **(h)** Z-score distribution of proteins categorized by the outlier status of the promoter that transcripts them.

To investigate the global relationship between accessibility outliers and gene expression, we compared the variations in promoter accessibility with variations in gene expression across all samples (Fig. 3f). We observed a significant correlation between the fold changes (*log*_2_(*FC)*) in accessibility at genes’ promoters and the respective expression levels of these genes (Spearman’s correlation coefficient=71%, *P* < 0. 001). The high correlation indicates that aberrations in promoter accessibility potentially influence the aberrations in gene expression.

Moreover, we demonstrated that gene expression outliers can be systematically predicted from the promoters, proximal, and distal enhancer accessibility (Fig. 3g). Ranking promoter outliers by their absolute log fold change (|log_2_(*FC)*|) to predict gene expression outliers achieve an auPRC of 11.7%. If the promoter is an outlier, there is a 44%chance (precision) that its gene is an expression outlier, and 24%of gene expression outliers have an outlier promoter (recall). Similarly, proximal enhancer outliers are highly predictive of gene expression outliers. Specifically, 27% (the recall at 21%precision) of gene expression outliers have at least one outlier proximal enhancer. Ranking genes based on the absolute log fold change of their outlier proximal enhancers achieves the performance of auPRC of 6.2%. We also ranked genes based on the maximum absolute fold change of accessibility outliers in 100 kilo-bp vicinity regardless of annotation of outliers and achieved 3%auPRC. We weighted the absolute log fold change of distal enhancers by ABC score^36^ and obtained a score for each gene (Methods). The score calculated from distal enhancer outliers is also a reliable predictor of gene expression outliers and achieves 5.2%auPRC. We trained an explainable boosting machine^37^ to predict gene expression outliers by combining features from promoters, proximal, and distal enhancers (Methods). The machine learning model achieved 21.9%auPRC. Altogether, these results show that chromatin accessibility outliers are predictive for gene expression outliers, and chromatin accessibility aberrations in promoters, proximal, or distal enhancers often translate into gene expression aberrations.

We focused on gene expression outliers with outlier promoters and investigated aberrations in their protein levels (Fig. 3h). We observed that the gene expression outliers with overly accessible promoters have higher protein levels (average Z-score = 2.2, *P* < 0. 001 based on the Mann– Whitney U test), and genes with under-accessible promoters have lower protein levels (average Z-score = 2.2, *P* < 0. 001 based on the Mann–Whitney U test). Overall, we present the biological significance of accessibility outliers on molecular phenotype by replicating outliers from multiple omics levels.

### A comparison of chromatin accessibility and gene expression reveals whether aberrations in molecular phenotype are linked to transcriptional or post-transcriptional regulation

The interplay between transcription and degradation rates determines RNA levels (Fig. 4a). Aberrant promoter or enhancer activity can lead to up or down-regulation of gene expression by altering transcription. Alternatively, alteration of post-transcriptional processes may affect RNA degradation, notably but not exclusively via non-sense-mediated decay (NMD) ^16^. Hence, the potential regulatory process associated with aberrant gene expression can be revealed by comparing accessibility outliers against gene expression outliers.

**Fig. 4:**
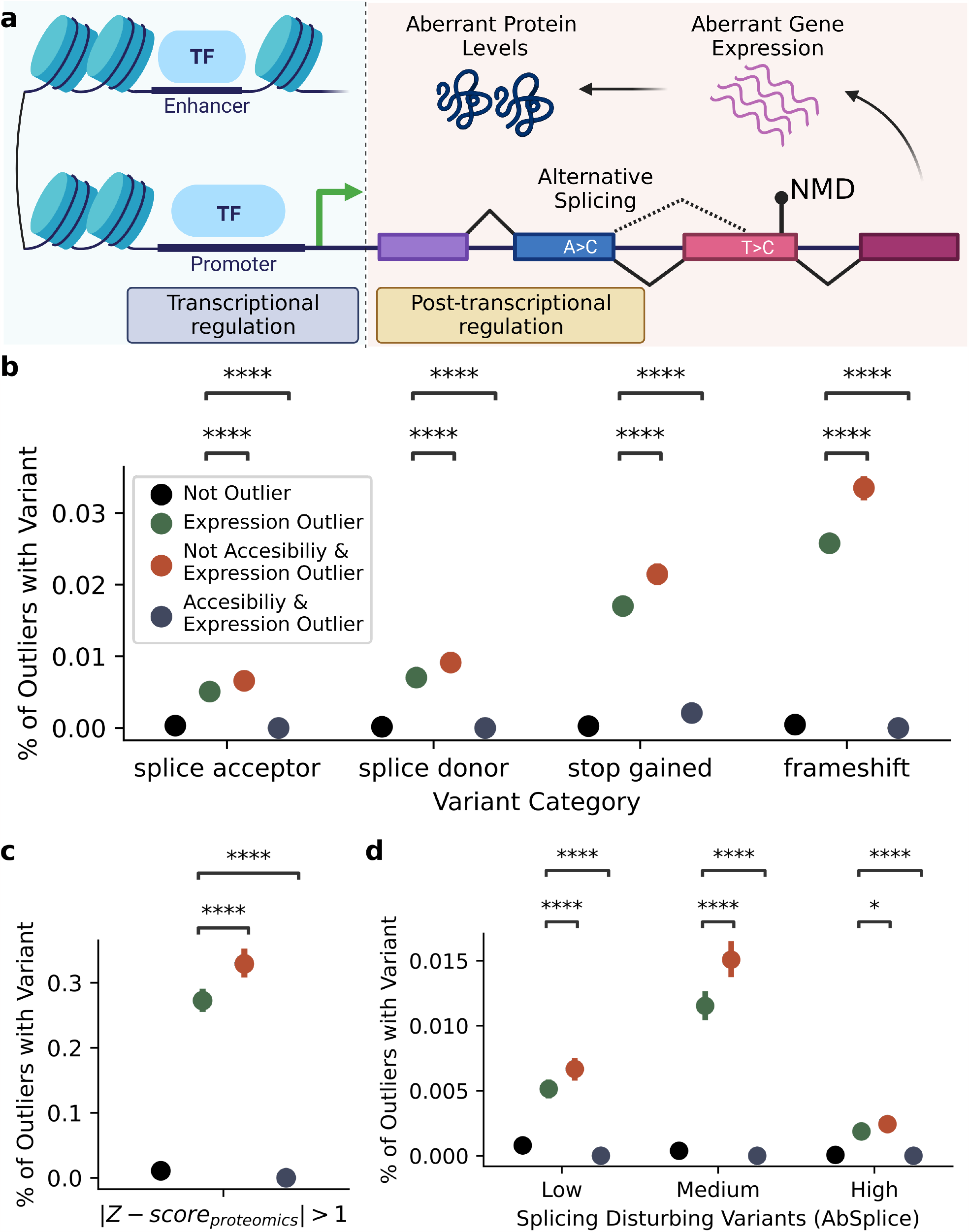
Overlap between accessibility outliers, gene expression outliers, and rare genetic variants by consequence. **(a)** Aberration in gene expression can result from dysregulation in transcriptional regulation, such as aberrant promoter or enhancer activity, or dysregulation in post-transcriptional regulation, such as splicing or nonsense-mediated decay. **(b)** Percentage of genes with potentially NMD-triggering rare genetic variants by outlier categories. Gene expression outliers without outlier promoters are enriched, and gene expression outliers with promoter outliers are depleted for variants in each category. **(c)** Percentage of protein with a missense or potentially NMD-triggering rare variants where protein levels are at least deviant by |*Z*-score| > 1 by outlier categories. **(d)** Percentage of genes containing splicing-disrupting variants predicted by AbSplice by outlier categories.

To demonstrate this point, we investigated the enrichment of rare and potentially NMD-triggering variants in different outlier categories. Fig. 4b presents the frequency of potentially NMD-triggering rare variants, such as splice acceptor, donor, nonsense, and frameshift variants, based on their SnpEff^38^ consequences by the outlier type of the affected gene. These variants were observed rarely and appeared in fewer than 0.1% of non-outlier genes. Compared to the baseline of all expression outliers, these variants are significantly more prevalent in gene expression outliers if their promoters are not accessibility outliers. The enrichment holds for splicing acceptor (n = 11, *P* < 0. 001 based on the Mann–Whitney U test calculated with bootstrapping) and splicing donor (n = 15, *P* < 0. 001), nonsense (n = 39, *P* < 0. 001), and frameshift variants (n = 52, *P* < 0. 001). Remarkably, only one of the gene expression outliers with aberrant promoter activity contains potentially NMD-triggering rare variants, indicating a significant depletion pattern.

Further investigation of both missense and potentially NMD-triggering rare variants in genes with aberrant promoter accessibility and protein levels presents a similar trend of depletion (Fig. 4c). 25.9% of expression outlier genes (*n* = 108) with aberrant protein levels (|Z-score| > 1) contain at least one of such variants. When genes with promoter outliers were excluded, the enrichment of variants rose to 3 0.7% (n = 91, *P* < 0. 001 based on the Mann–Whitney U test calculated with bootstrapping). The remaining expression outlier genes (*n* = 17) have promoters with aberrant accessibility, and none of these genes contain genetic variants with mentioned consequences. The substantial depletion of these variants in these genes (*P* < 0. 001) indicates that their aberrant protein levels are potentially linked to aberrant promoter accessibility rather than coding variants.

Aberrant splicing is another mechanism that can affect gene expression by resulting in aberrant RNA isoforms subject to NMD^39^. Thus, we further explored the impact of exonic or intronic splicing-disrupting variants prioritized by AbSplice^40^. We detected 40 gene expression outliers containing at least one splicing-disrupting variant across different thresholds (Fig. 4d). However, none of these genes has an outlier promoter regardless of the threshold choice. In contrast, subsetting gene expression outliers without aberrant promoters increases the prevalence of splicing disturbing variants for the subset.

The results underscore that comparing accessibility outliers against gene expression outliers and aberrant protein levels can identify whether aberration in the molecular phenotype is tied to transcriptional or post-transcriptional regulation.

### Analyzing outliers across multiple omics levels reveals potentially novel candidate genes for ALS

We reviewed the literature to understand the biological significance of outliers observed across multiple omics levels in ALS samples. Among the outlier genes that exhibited aberrations in both promoter accessibility and gene expression (Supplementary Table-1), 12 have previously been associated with ALS: *CDKL5, HIF1A, ABCA2, VPS4B, NOVA1, NRG1, NIPA1, BCL2, ALYREF, UBQLN2, IRAK4*, and *DDX3X*^1,5,7,11,41–47^. In some cases, variants have been associated with ALS or pathways involving these genes are dysregulated. While several proteomics measurements were missing due to the limitations of mass spectrometry^48^, three genes from these outliers (*VPS4B, ALYREF, DDX3X*) also displayed aberrant protein levels. For instance, we observed elevated expression and protein levels of *ALYREF*, in accordance with prior research (Extended Data Fig. 6) and knocking down an orthologue of *ALYREF* in an animal model reduces *TDP-43* induced toxicity^49^. Similarly, the *CDKL5* gene exhibits over-expression with an over-accessible promoter region (protein levels are unavailable). Suppressing *CDKL5* expression using a small molecule probe enhances the survival of human motor neurons under endoplasmic reticulum stress conditions^50^. Another outlier gene we identified, *HIF1A*, contributes to motor neuron degeneration through hypoxic stress, and prolonged survival observed in ALS mice suggests up-regulation of *HIF1A* as a potential therapeutic target^51^. Finally, *VPS4B* is pathologically increased in familial and sporadic ALS neuronal nuclei^52^. A closer examination of these identified outlier genes could reflect potential mechanisms involved in ALS and/or illuminate pathogenesis in subsets of ALS patients.

While some of the outlier genes are previously unreported as being associated with ALS, they play an important role in pathways involved in ALS. For example, the promoter of *LCMT1* is less accessible, and both its gene expression and protein level are downregulated in our dataset.Increased tau phosphorylation has been reported in ALS^53^ and down-regulation of *LCMT1*, in conjunction with the up-regulation of *HIF1A*, has been linked to tau hyperphosphorylation^5^. *DDX6* is another gene that is down-regulated across three omics levels (Extended Data Fig. 7) and is a loss-of-function intolerant gene^55^ (with a loss-of-function observed/expected upper bound fraction of LOEUF = 17%). Although the role of *DDX6* in ALS has not been documented, *DDX6* plays a critical role in RNA metabolism, particularly in the assembly of stress granules, a pathway dysregulated in ALS^56^. Furthermore, *DDX6* interacts with the ALS gene *ATXN2*, and a knockout of *DDX6* severely disrupts p-body formation^57^. In two ALS samples, we observed reduced promoter accessibility, gene expression, and protein levels of *NEDD4L* (Extended Data Fig. 8), another loss of function intolerant gene (LOEUF = 20%). *NEDD4L* is a direct substrate of *USP7* that regulates proteotoxicity in ALS^58^. Oxidative stress is implicated in neurodegeneration^59^, and up-regulation of *TXNL1* has been shown to reduce oxidative stress in neurological conditions^60,61^. Therefore, the pronounced down-regulation of promoter accessibility, gene expression, and protein levels of *TXNL1* observed in our study may be associated with increased oxidative stress (Extended Data Fig. 9). Vesicle transport is dysregulated by loss of function of ALS associated genes, such as *VAPB*^62^, *NEK1*^63^. *LMAN1*, a cargo receptor for the endoplasmic reticulum-Golgi transport^64^, is also involved in the trafficking of neuroreceptors^65^. The observed reduction in promoter accessibility, gene expression, and protein levels of *LMAN1* in two of our samples could be of relevance to ALS (Extended Data Fig. 10).

By cross-referencing genes with those associated with neurodegenerative disorders in OMIM^66^, we discovered six additional gene expression outliers (*GAN, EIF4A2, NARS1, HSD17B10, ERCC8, SLC25A46*) with aberrant promoter accessibility that are involved in both neurodegenerative and neurodevelopmental disorders. A notable example is *ERCC8*, involved in DNA damage repair and when mutated causes Cockayne Syndrome^67^, an early-onset degenerative condition^67^. *ERCC8* has also been identified as a comorbid factor in shared genetics between Parkinson’s disease and ALS^69^.

Overall, our multi-omics level analysis both reaffirms the involvement of known ALS genes and introduces novel genes that might play a role in the ALS disease pathways.

## Discussion

In this study, we introduced EpiOut, a computational toolbox for detecting and annotating chromatin accessibility outliers, which are characterized as large aberrations in a few regions specific to a single or few samples. We applied our proposed method to ATAC-seq data from ALS patients and clinically healthy individuals. Our methodology employs a negative binomial test for detecting outliers with statistical significance. The mean parameter of the negative binomial is fitted using a linear autoencoder (LR-AE), and the dispersion parameter is inferred based on observed and expected counts using maximum likelihood estimation. Controlling for both known and latent confounding factors is crucial to exclude outliers resulting from technical artifacts or confounding factors during outlier detection^14,30^. The proposed LR-AE is an effective statistical method for obtaining latent confounders from high-dimensional data. PCA, a specific case of an LR-AE, can also detect and correct for latent confounders better than alternative statistical methods^70^. However, PCA minimizes the Euclidean distance between measured and reconstructed counts; thus, it is suboptimal for discrete read counts of omics data with high dispersion and uncertainty due to low-coverage^71^. Thus, the negative binomial objective in our methodology is more apt for discrete read counts. Moreover, our software is more scalable than OUTRIDER and optimized for chromatin accessibility with a dimensionality of hundreds of thousands of genomic regions. While primarily designed for ATAC-seq, our toolbox is readily compatible with other accessibility assays, such as DNase-seq.

Our toolbox includes EpiAnnot, which annotates accessible regions for ChIP-seq marks. Using EpiAnnot, we found that outliers are enriched in functional regions, particularly promoters and active enhancers. Interestingly, outlier pairs tend to occur nearby, with many promoter outliers tied to active enhancer outliers within one million bp vicinity. This observation is supported by the relatively high Hi-C contact scores for outlier pairs in the vicinity, indicating potential interactions between these outliers.

By examining multiple omics levels, we found consistent replication of outliers. Accessibility outliers are associated with downstream biological processes such as gene expression and protein levels. In particular, a significant proportion of the gene expression outliers can be predicted from the aberrant accessibility of the promoter, proximal, and distal enhancer regions. Similarly, aberrant promoter activity is correlated with up and down-regulation of protein levels.

Analyzing the interplay between accessibility and gene expression outliers yields insight into whether aberration in gene expression originates from transcriptional regulation, such as increased synthesis rate via higher promoter activity, or post-transcriptional regulation, such as splicing or nonsense-mediated decay. We observed substantial depletion of NMD-triggering rare variants in gene expression outliers if promoters of these genes are an accessibility outlier and conversely observed enrichment of these variants if their promoter is not an outlier.

Our outlier detection methodology offers a novel avenue for studying rare diseases. We have successfully adapted the outlier detection approach to ATAC-seq data and underscored that chromatin accessibility is a beneficial complementary assay for rare disease diagnostics. Detected outliers in ALS samples are highly robust and consistently replicated across multiple omics levels. Many of these outliers are either known ALS genes or are involved in pathways implicated in ALS. Thus, the continued development and integration of the outlier detection approach with disease gene discovery methodologies may ultimately lead to a more comprehensive understanding of the genetic factors contributing to ALS. Such advancements could finally bridge the gap between the disease heritability and the known catalog of ALS disease genes.

## Method

### AnswerALS dataset

The multi-omics dataset, which includes ATAC-seq, RNA-seq, proteomics, and whole genome sequencing (WGS), for Amyotrophic Lateral Sclerosis (ALS) was downloaded from the Answer ALS portal (dataportal.answerals.org). The data contains 245 individuals diagnosed with ALS and 45 samples from clinically healthy controls. All samples had corresponding ATAC-seq and RNA-seq experiments. 253 samples also have whole genome sequencing (WGS) data. For the scope of our study, we restricted our analysis to the 253 samples that had paired data across three omics levels. Additionally, 204 of these samples have proteomics data. In the proteomics-related analysis, we subset and only used these samples with the proteomics data.

### Peak calling

We performed joint peak calling on ATAC-seq data to detect accessible regions across all samples using MACS2^28^. First, we merged bam files from every sample into a unified bam file utilizing SAMtools^72^, and subsequently filtered out reads with a mapping quality (MAPQ) below 10. Duplicate reads were retained after merging reads across samples. The default arguments of MACS2 were used, except for the duplicate read filter. ATAC-seq peaks contained in the narrow peak bed file generated by MACS2 were used in the downstream analysis.

### Read counting

We implemented an ultra-fast read counting algorithm for ATAC-seq described in Supplementary Algorithm-1. The counting algorithm is optimized based on two primary assumptions: ATAC-seq peaks are not overlapping and are separated by at least a gap longer than a read length; moreover, both peaks and reads are sorted by the genomic coordinates. We ensure the first assumption by jointly calling peaks as described above and collapsing any peaks closer to each other than the minimum gap distance (default 200 bp); thus, an ATAC-seq read never intersects with two peaks simultaneously. Since the bam file format is pre-sorted by genomic coordinates, ATAC-seq peaks are sorted to guarantee the second condition before the counting step. The counting algorithm creates two stacks of sorted peaks and reads, iterates over reads and peaks, and tracks the number of overlaps (Supplementary Algorithm-1). The resulting runtime complexity of the counting step is O(*r* + *p)* where r is the number of reads, and p is the number of peaks. In the preprocessing step, peaks are sorted, so the overall complexity is O(*p log*(*p*) + *r)*. However, since the number of reads is much larger than the number of peaks (r > p), the algorithm practically behaves in linear runtime in terms of the number of reads. The memory complexity of counting is linear in terms of the number of peaks (O(*p*)) because reads are fetched iteratively per chromosome by leveraging the indexable file format of BAM.

### Filtering peaks by replication rate

We applied two filters to the ATAC-seq peaks detected by MACS2 to ensure consistent peak replication across samples. First, we eliminated peaks with low coverage, specifically those with fewer than 100 reads across all samples. As a subsequent criterion, any peak must be supported by a minimum of 2 reads in at least 50% of all samples.

### Size factor normalization

To account for coverage differences across the ATAC-seq experiments, we performed size factor normalization on the read counts, a method initially proposed by DESeq2^31^. The size factor *s*_*i*_ for a sample *i* is defined as:

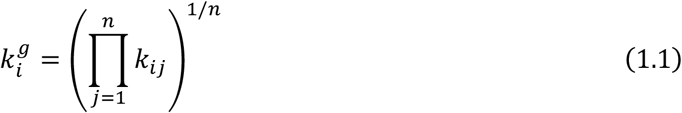

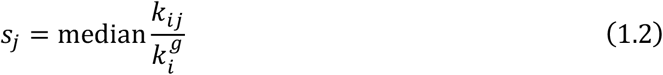

where *k*_*ij*_ represents the number of reads mapped to region *j* in sample *i* and 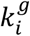 is the geometric mean of the reads across regions for sample *i*. The median of *k*_*ij*_ to 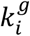 ratios is defined as the size factor *s*_*i*_ of sample *i*.

### Outlier detection

After normalizing the read counts of peaks with size factor normalization, we log-transformed the normalized counts and centered them around zero by subtracting the mean normalized counts:

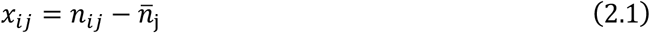

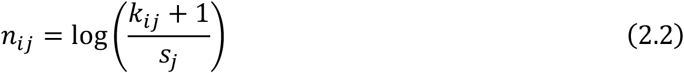

Autoencoder is applied on *x*_*ij*_ to calculate the expectation of normalized read counts 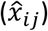. *f*_*e*_ is the encoder function of the autoencoder, which takes observed normalized counts (*x*_*ij*_) as input and calculates major covariates. Major covariates are concatenated with known confounders to obtain latent representation (h), which is decoded back to expected normalized counts 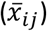 with the decoder function *f*_*d*_:

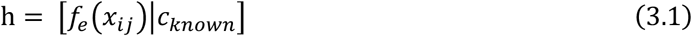

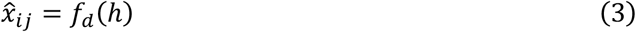

We used linear encoder and decoder functions. The encoder model uses a principal component analysis (PCA), where the encoder weights (*W*_*e*_) are the rotation matrix of PCA. The decoder weights (*W*_*d*_) are initialized with linear regression and further trained with the negative binomial objective:

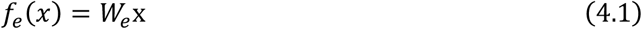

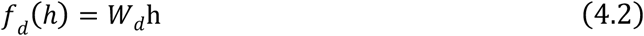

Log normalized counts (*x*_*ij*_) is transformed back to the original natural scale:

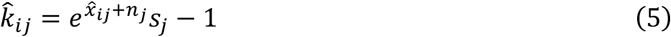

The dispersion parameter is optimized with maximum likelihood estimation (MLE) where the likelihood function:

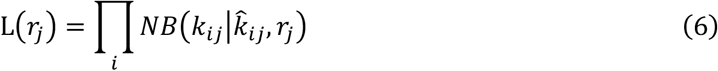

Fitting the dispersion parameter requires solving an independent convex problem for each peak. To solve a large number of independent convex problems quickly, we implemented a vectorized backtracking line search algorithm using TensorFlow. The dispersion parameter range is set between a lower bound of 0.01 and an upper bound of 1,000 to avoid numerical stability issues and overfitting.

P-value (*P*_*ij*_) of each peak and sample is calculated with a two-sided negative binomial test:

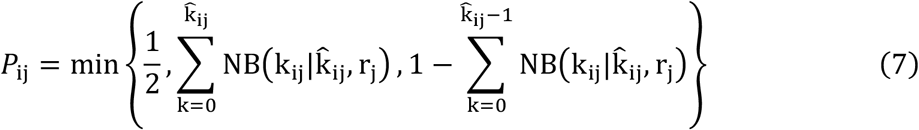

Finally, p-values are corrected for multiple testing corrections to control the false discovery rate with the Benjamini-Yekutieli method.

In addition to p-values, we report the log of fold-change or the log ratio of observed read counts to expected read counts calculated by:

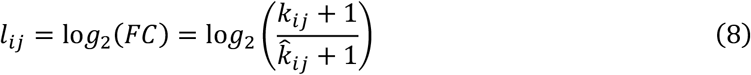

and Z-score based on fold changes defined as:

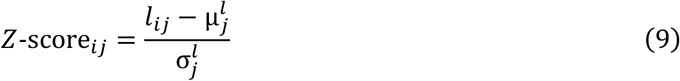

We consider peaks as outliers if their adjusted p-values are smaller than 5%(*P*_ad*j*_ < 0. 05), their absolute log fold changes are greater than 5 0%(|log_2_(FC)| > 0.5) and either their read counts (*k*_*ij*_) or expected read counts 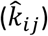 are at least 5 0.

### Injection of artificial outliers

In order to benchmark the performance of outlier detection methods, we conducted a simulation experiment using artificial outliers. We created an outlier mask *M*_*ij*_ for sample-peak pairs. A sample-region pair is categorized as either an over-accessibility or under-accessibility outlier with a probability of 0.01%:

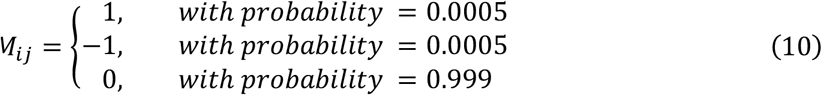

Log-normalized counts are updated using an artificial outlier mask:

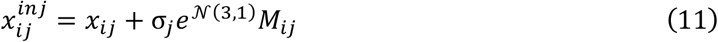

where the deviation in accessibility outlier is simulated by scaling the standard deviation of the peak (σ_*j*_) using a value sampled from a log-normal distribution parameterized by a mean of 3 and a standard deviation of 1. Eq. 5 transforms log-transformed normalized injected counts 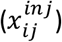 back to counts in the natural count scale 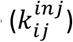.

Outlier detection methods are evaluated by benchmarking their capability to predict the outlier mask (*M*_*ij*_) from the injected counts 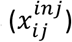. The primary benchmark metric for this evaluation is the area under the precision-recall curve (auPRC). We generated ten outlier masks for testing and one additional mask for the validation set. Hyperparameter tuning was performed on the validation set to identify the optimal bottleneck size of autoencoder-based models. The evaluation procedure was executed ten times on the test folds, and the average auPRC performance and its standard deviation are reported.

We evaluated four methods: naive negative-binomial test, PCA, OUTRIDER, and EpiOut. During the precision-recall calculation, predictions for each method were primarily ranked using p-values, except for PCA, which was ranked by its Z-score. The performance of each method at a specific cutoff (either at the adjusted p-values of 0.05 or the absolute Z-score of 2) is indicated on the precision-recall curve.

For the naive negative binomial test, we averaged counts across samples per peak to obtain the mean parameter of the negative binomial distribution, 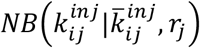. The dispersion is estimated with MLE (Eq. 6).

To evaluate the PCA model, we used PCA to estimate the expected read counts, 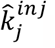. These expected read counts are the reconstructed read counts of PCA. The number of principal components retained was determined through hyperparameter tuning on the validation set.

Predictions were subsequently ranked based on the Z-score, as formulated in Eq. 9, using the expected counts from PCA. We did not compute dispersion or perform a negative binomial test.

We ran OUTRIDER and EpiOut with their default parameters. The bottleneck size of all autoencoder-based methods was selected based on the hyperparameter tuning on the validation set.

### Cross-correlation between sample

We used Pearson’s correlation to calculate the cross-correlation between samples. The raw cross-correlations were obtained using observed counts (k_*ij*_) without any transformation. The cross-correlation after correction was calculated using corrected counts:

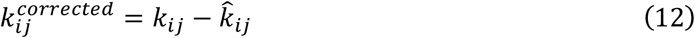

### Runtime benchmark

To measure runtime to call outliers with OUTRIDER and EpiOut, we subset peaks into groups of 10000, 25000, 50000, and 100000 and calculated runtime for each input size. 8 CPU cores are used during the benchmark. The analysis is repeated 10 times. The average runtime and standard deviation of runtime were reported. We ran each tool with bottleneck sizes of 5, 10, 50, and 100 in each iteration to avoid runtime differences resulting from hyperparameter tuning. We reported the total runtime across different bottleneck sizes.

### Functional annotation of accessible regions with ChIP-seq

We introduced EpiAnnot, a software tool for functional annotation of accessible genomic regions using ChIP-seq marks. If a specific ChIP-seq for a cell line or tissue is present in the Roadmap Epigenomics^32^ or ENCODE^33^ databases, EpiAnnot can retrieve ChIP-seq data from these public data sources. Additionally, users can annotate their accessible regions using their own custom ChIP-seq data.

EpiAnnot can also attribute accessible regions to genomic features using a gene transfer format (GTF) file. For instance, it annotates accessible regions as 5’ UTR, TSS, etc. In scenarios where an accessible region overlaps with the H3K4me3 histone ChIP-seq mark and TSS or 5’ UTR of a gene, EpiAnnot designates these regions as promoters. Conversely, regions intersecting with the H3K4me1 histone ChIP-seq mark are labeled as enhancers. When the histone mark is available, enhancers overlapping with H3K27ac are categorized as active; otherwise, they are poised. EpiAnnot also annotates enhancers as proximal or distal based on their distance from genes. Specifically, enhancers within a gene body or located up to 10,000 bp upstream or 2,000 bp downstream from a gene are tagged as proximal. All others are marked as distal enhancers.

By using EpiAnnot with H3K4me3, H3K27ac, and H3K4me1 histone ChIP-seq marks from in vitro differentiated motor neurons from ALS and clinically healthy samples^34^, we annotated accessible regions from ATAC-seq experiments and used the GTF file of GENCODE v38. The histone ChIP-seq data was downloaded from ENCODE.

### Enrichment of outlier pairs in the proximity

To investigate the potential interaction between nearby outlier pairs, we quantified the pairs of accessible regions within predefined distances, irrespective of the outlier status of regions. Then, we applied Fisher’s exact test to calculate the enrichment of outlier pairs, using a contingency table structured with the outlier statuses of accessible regions in the pair. We repeated this analysis for 10,000, 100,000, 500,000, and 1,000,000 bp distances. The odds ratio was also calculated to highlight the likelihood of the region being an outlier, given the outlier status of the nearby region.

### Annotation of chromatin interactions with Hi-C

To delve deeper into the interactions between outliers, we annotated their interactions based on Hi-C contacts using EpiAnnot. We downloaded public Hi-C data from motor neurons differentiated from iPSCs^34^ from ENCODE and computed contact scores based on the number of Hi-C reads between accessible regions using EpiAnnot. For the interaction analysis, the genome is partitioned into bins of 5,000 bp. The interaction between a first source region (*s*) and a target second region (*t*) is determined by the highest Hi-C contact score between these bins or their immediate neighbors:

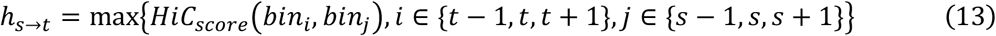

We fitted a power regression to estimate Hi-C contact scores between pairs of accessible regions, using the outlier status of the pairs while controlling for distance. P-values for the regression coefficients were calculated using a t-test.

We utilized EpiAnnot to compute the ABC score, which signifies potential interactions between regions informed by Hi-C contact scores. The ABC score is defined as:

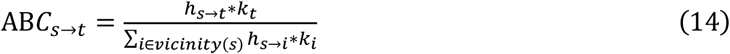

where the numerator represents the Hi-C contact score between regions multiplied by the accessibility of the target region (*k*_t_ = ∑_*i*_ *k*_*ij*_) and the denominator normalizes this value for all region pairs associated with the source region.

### Aberrant gene expression prediction from chromatin accessibility

Gene expression outliers were called with OUTRIDER using the DROP pipeline. If genes were deviant by least an absolute log-fold change of 30% (|log_2_(FC)| > 0.3), and their adjusted p-values based on the negative binomial test were smaller than 5% (*P*_ad*j*_ < 0. 05), they were considered expression outliers. These gene expression outliers constituted the ground truth labels in the benchmark. In order to predict gene expression outliers from accessibility outliers, we trained an explainable boosting machine (EBM). The EBM model was trained and tested with 5-fold cross-validation. The model aggregates features from promoters and enhancers as input to predict the probability of a gene being categorized as an expression outlier. For predicting a gene’s outlier status, the model considers the following features: log fold change, p-value, and outlier status of the promoter, maximum absolute log fold change of the proximal enhancer outliers, and the ABC-score weighted absolute log fold change of the distal enhancer outliers. The score for the weighted distal enhancer is computed as follows:

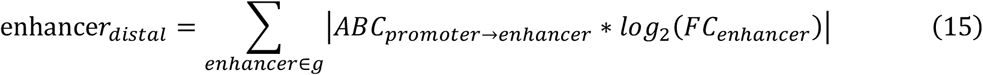

### Depletion of rare variants with certain consequences in genes with promoter outlier

We assessed the depletion of potentially NMD-triggering rare variants in the gene expression outliers where promoters of these genes are also outliers. Variants are annotated with SnpEff v4.3^38^ according to their consequences for genes. Variants with splice acceptor, donor, nonsense, and frameshift consequences were considered as potentially NMD-triggering. Only rare variants with a frequency of 0.1% were used in the analysis. Variant frequencies are downloaded from the gnomAD database^55^. Consequence categories used for depletion analysis include stop codon, frameshift, acceptor, donor site dinucleotides, and missense variants. AbSplice was used to classify splicing-disrupting exonic or intronic variants into high, medium, or low-impact categories, using thresholds of 0.2, 0.1, and 0.05, respectively, as advised by the authors. We determined the proportion of genes affected by each variant category based on the outlier status of the genes and their associated promoters. We repeated the enrichment analysis 100 times by randomly subsetting genes in each iteration for bootstrapping. We used the Mann–Whitney U test to measure statistical significance based on the proportions calculated with bootstrapping. Moreover, we analyzed proteomics data to quantify the proportion of proteins impacted by NMD-triggering or missense variants. Z-scores for the proteomics data were obtained with PROTRIDER.

### Curation of genes associated with neurodegenerative diseases

We manually curated a list of ALS genes from literature^1,11,42–47,49–51,73–82^ and ALSOD^41^. The curated list is available at 10.5281/zenodo.8331545. Moreover, we queried the OMIM^66^ database using “neurodegenerative” and “neurodegeneration” keywords, filtered the retrieved entries based on neurologic clinical synopsis, and generated a list of genes associated with neurodegenerative diseases via the REST API. The genes were further annotated for their loss of function intolerance obtained from gnomAD^55^. Genes are considered loss of function intolerant if their loss-of-function observed/expected upper bound fraction (LOEUF) is below 35%.

### Source Code

All the analyses in the paper are implemented in the reproducible snakemake^83^ format available at github.com/uci-cbcl/ALS-accessibility-outliers-paper. Epiout, the Python package for outlier detection and annotation, is available at github.com/uci-cbcl/EpiOut.

## Acknowledgments

This project is funded by a grant from the Robert Packard Center for ALS Research at Johns Hopkins and grants from the National Science Foundation [IIS-1715017, DMS-1763272]. Data used in the preparation of this article were obtained from the ANSWER ALS Data Portal (AALS-01184). For up-to-date information on the study, visit https://dataportal.answerals.org.

## References

1. Hardiman, O. et al. Amyotrophic lateral sclerosis. Nat. Rev. Dis. Primer 3, 1–19 (2017).

2. Longinetti, E. & Fang, F. Epidemiology of amyotrophic lateral sclerosis: an update of recent literature. Curr. Opin. Neurol. 32, 771–776 (2019).

3. Masrori, P. & Van Damme, P. Amyotrophic lateral sclerosis: a clinical review. Eur. J. Neurol. 27, 1918–1929 (2020).

4. Al-Chalabi, A. et al. An estimate of amyotrophic lateral sclerosis heritability using twin data. J. Neurol. Neurosurg. Psychiatry 81, 1324–1326 (2010).

5. Grassano, M. et al. Systematic evaluation of genetic mutations in ALS: a population-based study.J. Neurol. Neurosurg. Psychiatry 93, 1190–1193 (2022).

6. Renton, A. E., Chiò, A. & Traynor, B. J. State of play in amyotrophic lateral sclerosis genetics. Nat. Neurosci. 17, 17–23 (2014).

7. Akçimen, F. et al. Amyotrophic lateral sclerosis: translating genetic discoveries into therapies. Nat. Rev. Genet. 24, 642–658 (2023).

8. Van Rheenen, W. et al. Project MinE: study design and pilot analyses of a large-scale wholegenome sequencing study in amyotrophic lateral sclerosis. Eur. J. Hum. Genet. 26, 1537–1546 (2018).

9. Baxi, E. G. et al. Answer ALS, a large-scale resource for sporadic and familial ALS combining clinical and multi-omics data from induced pluripotent cell lines. Nat. Neurosci. 25, 226–237 (2022).

10. van Rheenen, W. et al. Common and rare variant association analyses in amyotrophic lateral sclerosis identify 15 risk loci with distinct genetic architectures and neuron-specific biology. Nat. Genet. 53, 1636–1648 (2021).

11. Gregory, J. M., Fagegaltier, D., Phatnani, H. & Harms, M. B. Genetics of Amyotrophic Lateral Sclerosis. Curr. Genet. Med. Rep. 8, 121–131 (2020).

12. NeuroLINCS Consortium et al. An integrated multi-omic analysis of iPSC-derived motor neurons from C9ORF72 ALS patients. iScience 24, 103221 (2021).

13. Frésard, L. et al. Identification of rare-disease genes using blood transcriptome sequencing and large control cohorts. Nat. Med. 25, 911–919 (2019).

14. Mertes, C. et al. Detection of aberrant splicing events in RNA-seq data using FRASER. Nat. Commun. 12, 529 (2021).

15. Yépez, V. A. et al. Detection of aberrant gene expression events in RNA sequencing data. Nat. Protoc. 16, 1276–1296 (2021).

16. Li, X. et al. The impact of rare variation on gene expression across tissues. Nature 550, 239–243 (2017).

17. Marwaha, S., Knowles, J. W. & Ashley, E. A. A guide for the diagnosis of rare and undiagnosed disease: beyond the exome. Genome Med. 14, 23 (2022).

18. Jenkinson, G. et al. LeafCutterMD: an algorithm for outlier splicing detection in rare diseases. Bioinformatics 36, 4609–4615 (2020).

19. Kremer, L. S. et al. Genetic diagnosis of Mendelian disorders via RNA sequencing. Nat. Commun. 8, 15824 (2017).

20. Kopajtich, R. et al. Integration of proteomics with genomics and transcriptomics increases the diagnostic rate of Mendelian disorders. 2021.03.09.21253187 Preprint at 10.1101/2021.03.09.21253187 (2021).

21. Li, T. et al. The functional impact of rare variation across the regulatory cascade. Cell Genomics 100401 (2023) doi:10.1016/j.xgen.2023.100401.

22. Klemm, S. L., Shipony, Z. & Greenleaf, W. J. Chromatin accessibility and the regulatory epigenome. Nat. Rev. Genet. 20, 207–220 (2019).

23. Liu, Q. et al. Chromatin accessibility landscapes of skin cells in systemic sclerosis nominate dendritic cells in disease pathogenesis. Nat. Commun. 11, 5843 (2020).

24. Turner, A. W. et al. Single-nucleus chromatin accessibility profiling highlights regulatory mechanisms of coronary artery disease risk. Nat. Genet. 54, 804–816 (2022).

25. Morabito, S. et al. Single-nucleus chromatin accessibility and transcriptomic characterization of Alzheimer’s disease. Nat. Genet. 53, 1143–1155 (2021).

26. Yan, F., Powell, D. R., Curtis, D. J. & Wong, N. C. From reads to insight: a hitchhiker’s guide to ATAC-seq data analysis. Genome Biol. 21, 22 (2020).

27. Abadi, M. et al. TensorFlow: Large-Scale Machine Learning on Heterogeneous Distributed Systems.

28. Zhang, Y. et al. Model-based analysis of ChIP-Seq (MACS). Genome Biol. 9, R137 (2008).

29. Stovner, E. B. & Sætrom, P. PyRanges: efficient comparison of genomic intervals in Python. Bioinforma. Oxf. Engl. 36, 918–919 (2020).

30. Brechtmann, F. et al. OUTRIDER: A Statistical Method for Detecting Aberrantly Expressed Genes in RNA Sequencing Data. Am. J. Hum. Genet. 103, 907–917 (2018).

31. Love, M. I., Huber, W. & Anders, S. Moderated estimation of fold change and dispersion for RNA-seq data with DESeq2. Genome Biol. 15, 550 (2014).

32. Kundaje, A. et al. Integrative analysis of 111 reference human epigenomes. Nature 518, 317–330 (2015).

33. Dunham, I. et al. An integrated encyclopedia of DNA elements in the human genome. Nature 489, 57–74 (2012).

34. Zhang, S. et al. Genome-wide identification of the genetic basis of amyotrophic lateral sclerosis. Neuron 110, 992–1008.e11 (2022).

35. Heintzman, N. D. et al. Distinct and predictive chromatin signatures of transcriptional promoters and enhancers in the human genome. Nat. Genet. 39, 311–318 (2007).

36. Fulco, C. P. et al. Activity-by-contact model of enhancer–promoter regulation from thousands of CRISPR perturbations. Nat. Genet. 51, 1664–1669 (2019).

37. Nori, H., Jenkins, S., Koch, P. & Caruana, R. InterpretML: A Unified Framework for Machine Learning Interpretability. Preprint at 10.48550/arXiv.1909.09223 (2019).

38. Cingolani, P. et al. A program for annotating and predicting the effects of single nucleotide polymorphisms, SnpEff. Fly (Austin) 6, 80–92 (2012).

39. Daguenet, E., Dujardin, G. & Valcárcel, J. The pathogenicity of splicing defects: mechanistic insights into pre-mRNA processing inform novel therapeutic approaches. EMBO Rep. 16, 1640–1655 (2015).

40. Çelik, M. H. et al. Aberrant splicing prediction across human tissues. Preprint at 10.1101/2022.06.13.495326 (2022).

41. Abel, O., Powell, J. F., Andersen, P. M. & Al-Chalabi, A. ALSoD: A user-friendly online bioinformatics tool for amyotrophic lateral sclerosis genetics. Hum. Mutat. 33, 1345–1351 (2012).

42. Pecoraro, V. et al. The NGS technology for the identification of genes associated with the ALS. A systematic review. Eur. J. Clin. Invest. 50, e13228 (2020).

43. Cheng, W. et al. CRISPR-Cas9 Screens Identify the RNA Helicase DDX3X as a Repressor of C9ORF72 (GGGGCC)n Repeat-Associated Non-AUG Translation. Neuron 104, 885–898.e8 (2019).

44. Krach, F. et al. Aberrant NOVA1 function disrupts alternative splicing in early stages of amyotrophic lateral sclerosis. Acta Neuropathol. (Berl.) 144, 413–435 (2022).

45. Mòdol-Caballero, G. et al. Therapeutic Role of Neuregulin 1 Type III in SOD1-Linked Amyotrophic Lateral Sclerosis. Neurotherapeutics 17, 1048–1060 (2020).

46. Blauw, H. M. et al. NIPA1 polyalanine repeat expansions are associated with amyotrophic lateral sclerosis. Hum. Mol. Genet. 21, 2497–2502 (2012).

47. Tazelaar, G. H. P. et al. Association of NIPA1 repeat expansions with amyotrophic lateral sclerosis in a large international cohort. Neurobiol. Aging 74, 234.e9-234.e15 (2019).

48. Poulos, R. C. et al. Strategies to enable large-scale proteomics for reproducible research. Nat. Commun. 11, 3793 (2020).

49. Berson, A. et al. Drosophila Ref1/ALYREF regulates transcription and toxicity associated with ALS/FTD disease etiologies. Acta Neuropathol. Commun. 7, 65 (2019).

50. Ong, H. W. et al. Discovery of a Potent and Selective CDKL5/GSK3 Chemical Probe That Is Neuroprotective. ACS Chem. Neurosci. 14, 1672–1685 (2023).

51. Nomura, E. et al. Imaging Hypoxic Stress and the Treatment of Amyotrophic Lateral Sclerosis with Dimethyloxalylglycine in a Mice Model. Neuroscience 415, 31–43 (2019).

52. Coyne, A. N. & Rothstein, J. D. The ESCRT-III protein VPS4, but not CHMP4B or CHMP2B, is pathologically increased in familial and sporadic ALS neuronal nuclei. Acta Neuropathol. Commun. 9, 127 (2021).

53. Stevens, C. H., Guthrie, N. J., van Roijen, M., Halliday, G. M. & Ooi, L. Increased Tau Phosphorylation in Motor Neurons From Clinically Pure Sporadic Amyotrophic Lateral Sclerosis Patients. J. Neuropathol. Exp. Neurol. 78, 605–614 (2019).

54. Lei, L. et al. HIF-1α Causes LCMT1/PP2A Deficiency and Mediates Tau Hyperphosphorylation and Cognitive Dysfunction during Chronic Hypoxia. Int. J. Mol. Sci. 23, 16140 (2022).

55. Karczewski, K. J. et al. The mutational constraint spectrum quantified from variation in 141,456 humans. Nature 581, 434–443 (2020).

56. Dudman, J. & Qi, X. Stress Granule Dysregulation in Amyotrophic Lateral Sclerosis. Front. Cell. Neurosci. 14, 598517 (2020).

57. Becker, L. A. et al. Therapeutic reduction of ataxin-2 extends lifespan and reduces pathology in TDP-43 mice. Nature 544, 367–371 (2017).

58. Zhang, T., Periz, G., Lu, Y.-N. & Wang, J. USP7 regulates ALS-associated proteotoxicity and quality control through the NEDD4L–SMAD pathway. Proc. Natl. Acad. Sci. 117, 28114–28125 (2020).

59. Hemerková, P. & Vališ, M. Role of Oxidative Stress in the Pathogenesis of Amyotrophic Lateral Sclerosis: Antioxidant Metalloenzymes and Therapeutic Strategies. Biomolecules 11, 437 (2021).

60. Zhao, J.-M. & Qi, T.-G. The role of TXNL1 in disease: treatment strategies for cancer and diseases with oxidative stress. Mol. Biol. Rep. 48, 2929–2934 (2021).

61. Yu, J.-T. et al. Up-regulation of antioxidative proteins TRX1, TXNL1 and TXNRD1 in the cortex of PTZ kindling seizure model mice. PLOS ONE 14, e0210670 (2019).

62. Tran, D., Chalhoub, A., Schooley, A., Zhang, W. & Ngsee, J. K. A mutation in VAPB that causes amyotrophic lateral sclerosis also causes a nuclear envelope defect. J. Cell Sci. 125, 2831–2836 (2012).

63. Mann, J. R. et al. Loss of function of the ALS-associated NEK1 kinase disrupts microtubule homeostasis and nuclear import. Sci. Adv. 9, eadi5548 (2023).

64. Zhang, Y. et al. LMAN1–MCFD2 complex is a cargo receptor for the ER-Golgi transport of α1antitrypsin. Biochem. J. 479, 839–855 (2022).

65. Fu, Y.-L., Zhang, B. & Mu, T.-W. LMAN1 (ERGIC-53) promotes trafficking of neuroreceptors. Biochem. Biophys. Res. Commun. 511, 356–362 (2019).

66. Hamosh, A., Scott, A. F., Amberger, J. S., Bocchini, C. A. & McKusick, V. A. Online Mendelian Inheritance in Man (OMIM), a knowledgebase of human genes and genetic disorders. Nucleic Acids Res. 33, D514–D517 (2005).

67. Taghdiri, M. et al. A Novel Mutation in ERCC8 Gene Causing Cockayne Syndrome. Front. Pediatr. 5, (2017).

68. Weidenheim, K. M., Dickson, D. W. & Rapin, I. Neuropathology of Cockayne syndrome: Evidence for impaired development, premature aging, and neurodegeneration. Mech. Ageing Dev. 130, 619–636 (2009).

69. Tian, Y. et al. Shared Genetics and Comorbid Genes of Amyotrophic Lateral Sclerosis and Parkinson’s Disease. Mov. Disord. n/a,.

70. Zhou, H. J., Li, L., Li, Y., Li, W. & Li, J. J. PCA outperforms popular hidden variable inference methods for molecular QTL mapping. Genome Biol. 23, 210 (2022).

71. Townes, F. W., Hicks, S. C., Aryee, M. J. & Irizarry, R. A. Feature selection and dimension reduction for single-cell RNA-Seq based on a multinomial model. Genome Biol. 20, 295 (2019).

72. Li, H. et al. The Sequence Alignment/Map format and SAMtools. Bioinformatics 25, 2078–2079 (2009).

73. Steinberg, K. M., Yu, B., Koboldt, D. C., Mardis, E. R. & Pamphlett, R. Exome sequencing of case-unaffected-parents trios reveals recessive and de novo genetic variants in sporadic ALS. Sci. Rep. 5, 9124 (2015).

74. Mishra, P. S. et al. Transmission of ALS pathogenesis by the cerebrospinal fluid. Acta Neuropathol. Commun. 8, 65 (2020).

75. Rotem, N. et al. ALS Along the Axons – Expression of Coding and Noncoding RNA Differs in Axons of ALS models. Sci. Rep. 7, 44500 (2017).

76. Hetz, C. et al. The proapoptotic BCL-2 family member BIM mediates motoneuron loss in a model of amyotrophic lateral sclerosis. Cell Death Differ. 14, 1386–1389 (2007).

77. Vukosavic, S., Dubois-Dauphin, M., Romero, N. & Przedborski, S. Bax and Bcl-2 Interaction in a Transgenic Mouse Model of Familial Amyotrophic Lateral Sclerosis. J. Neurochem. 73, 2460–2468 (2002).

78. Pasinelli, P. et al. Amyotrophic Lateral Sclerosis-Associated SOD1 Mutant Proteins Bind and Aggregate with Bcl-2 in Spinal Cord Mitochondria. Neuron 43, 19–30 (2004).

79. Yuan, Y. et al. Identification of GGC repeat expansion in the NOTCH2NLC gene in amyotrophic lateral sclerosis. Neurology 95, e3394–e3405 (2020).

80. Song, F., Chiang, P., Wang, J., Ravits, J. & Loeb, J. A. Aberrant Neuregulin 1 Signaling in Amyotrophic Lateral Sclerosis: J. Neuropathol. Exp. Neurol. 71, 104–115 (2012).

81. Schwenk, B. M. et al. TDP-43 loss of function inhibits endosomal trafficking and alters trophic signaling in neurons. EMBO J. 35, 2350–2370 (2016).

82. Nardo, G. et al. Amyotrophic Lateral Sclerosis Multiprotein Biomarkers in Peripheral Blood Mononuclear Cells. PLoS ONE 6, e25545 (2011).

83. Köster, J. & Rahmann, S. Snakemake--a scalable bioinformatics workflow engine. Bioinforma. Oxf. Engl. 28, 2520–2522 (2012).

