## Supplementary Material for "Identifying dysregulated regions in amyotrophic lateral sclerosis through chromatin accessibility outliers"

#### Extended Data

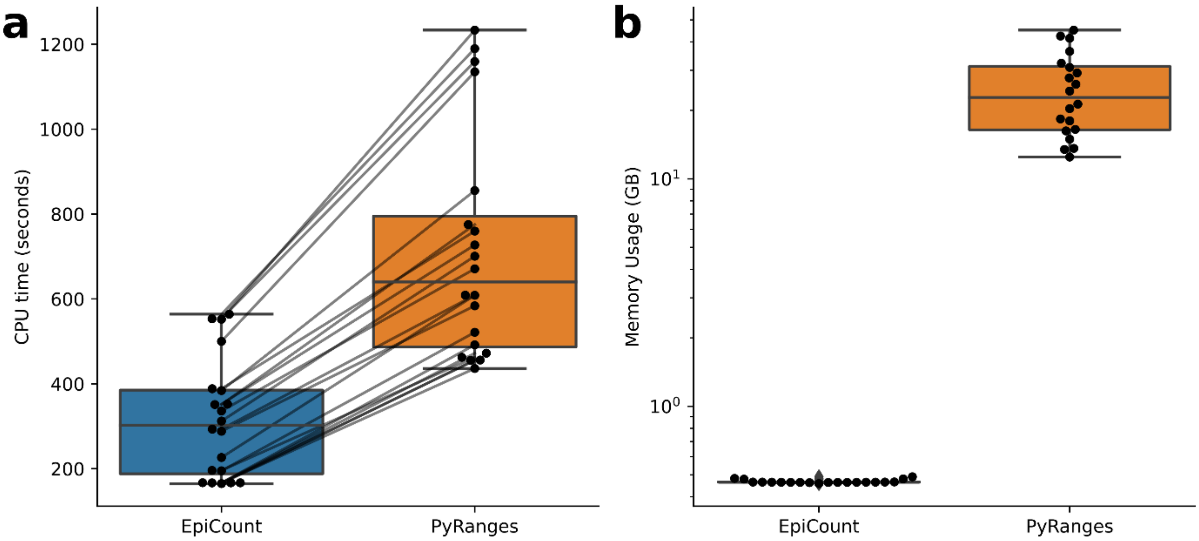


**Extended Data Fig. 1: Runtime of counting step.** Bam and bed files for 20 ATAC-seq experiments were downloaded from ENCODE. Each bam file has $142 \pm 48$ million ATAC-seq reads. Overlapping accessible regions across bed files were collapsed, leading to 365,090 accessible regions. The counting was performed with these accessible regions and bam files using EpiCount and PyRanges. **(a)** CPU time of EpiCount and PyRanges. EpiCount is twice as fast as PyRanges. The mean runtime of EpiOut per bam file is $316 \pm 139$ seconds; meanwhile, the runtime of PyRanges is $715 \pm 267$ seconds. Epiout counts one million reads per $2.1 \pm 0.2$ seconds, while PyRanges counts one million reads in $4.9 \pm 0.2$ seconds. **(b)** Epiout has a smaller memory footprint than PyRanges. PyRanges loads the entire bam file to memory while EpiCount performs stream counting by only loading a read per iteration; thus, EpiCount (mean memory consumption $0.46 \pm 0.008$ GB) consumes 2.1% of PyRanges' memory footprint (mean memory consumption $20.5 \pm 10$ GB). Extensive memory usage of PyRanges limits the parallelization across samples.

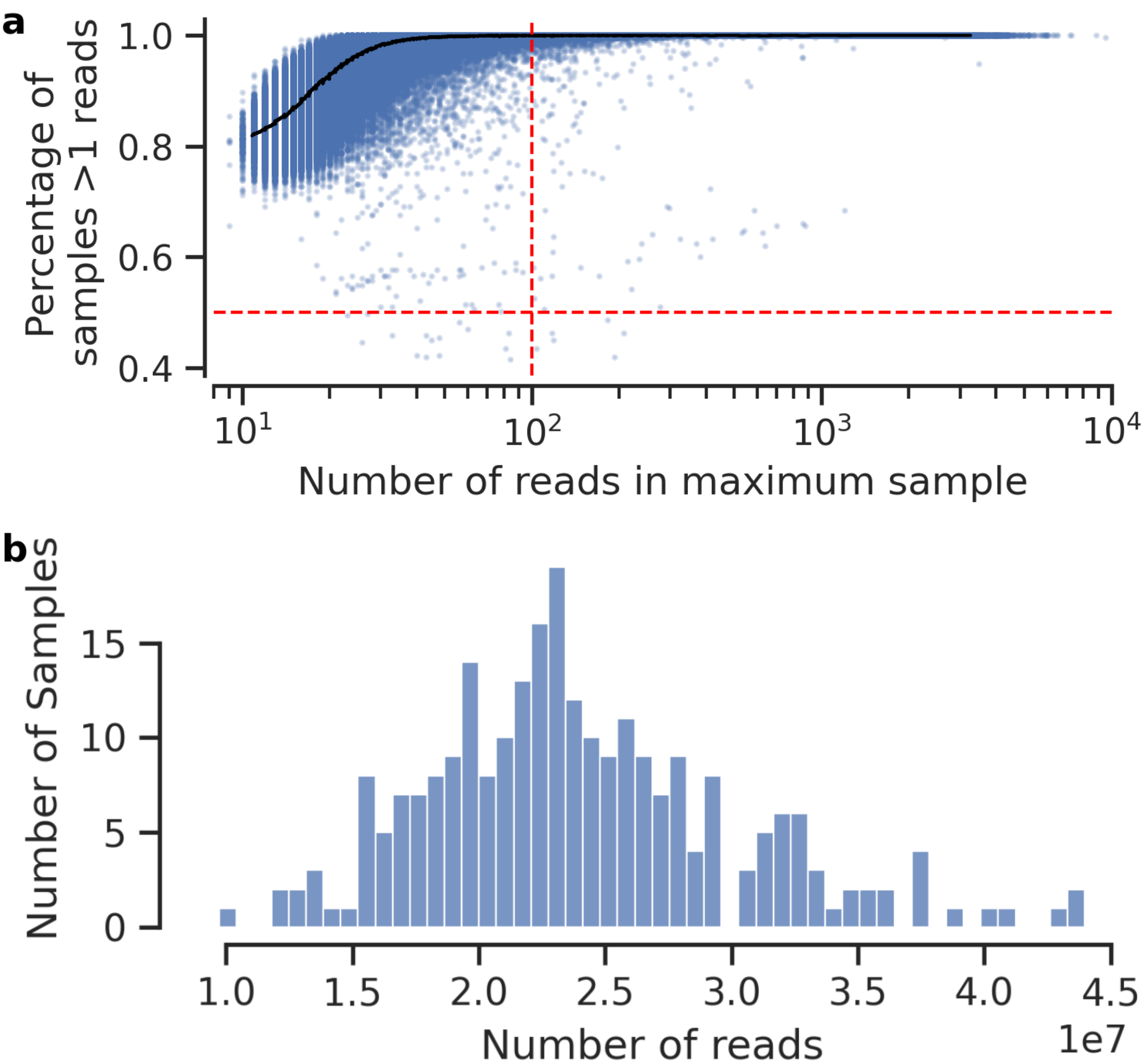


**Extended Data Fig. 2: Read coverage and replication rate statistics. (a)** Read coverage distribution across samples. Notable differences in read coverage are observed among the samples, highlighting the importance of size factor normalization to adjust for coverage disparity. **(b)** Replication rate of accessible regions throughout samples. In the scatterplot, each dot signifies an accessible region for a sample. The x-axis denotes the read counts supporting the accessible region in the sample with the highest read count. Filters, represented by the red lines, were implemented to ensure an accessible region is supported by at least 100 reads in one sample and is replicated in at least 50% of the samples by at least 2 reads.

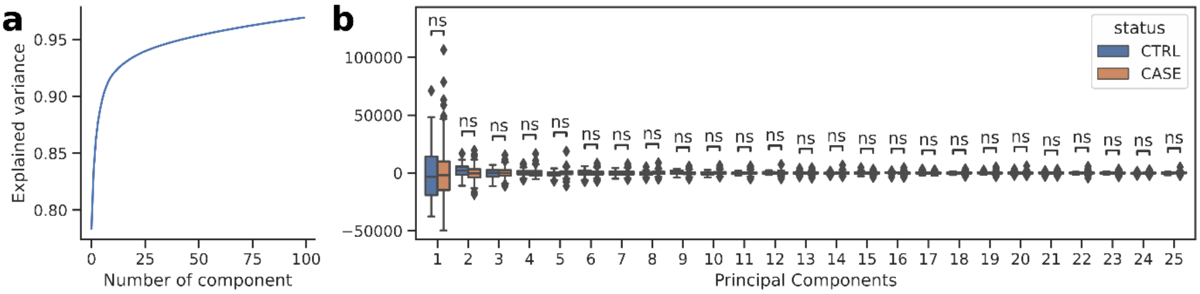


**Extended Data Fig. 3: The major covariates in the DNA accessibility data. (a)** Cumulative explained variance by top principal components. The top 25 principle components (representing the major latent confounding factors) explain $\sim 94\%$ of the accessibility covariation between samples. **(b)** The distribution of top principle components by the disease status. None of the top principal components are associated with the disease status based on the Mann–Whitney U test. P-values are corrected for multiple testing with the Benjamini-Yekutieli false-discovery rate (FDR) method.

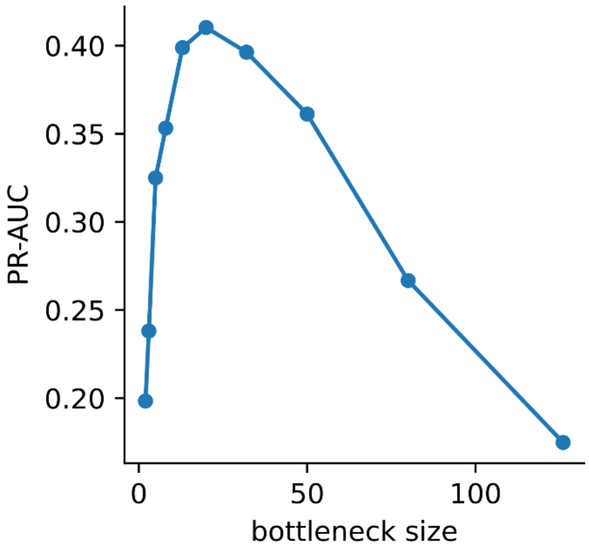


**Extended Data Fig. 4: Optimal bottleneck size choice with hyperparameter tuning.** EpiOut detects optimal bottleneck size with artificial outlier injection and hyperparameter tuning. The x-axis indicates the bottleneck size of LR-AE in the EpiOut model, and the y-axis shows the auPRC performance for the artificial outlier prediction.

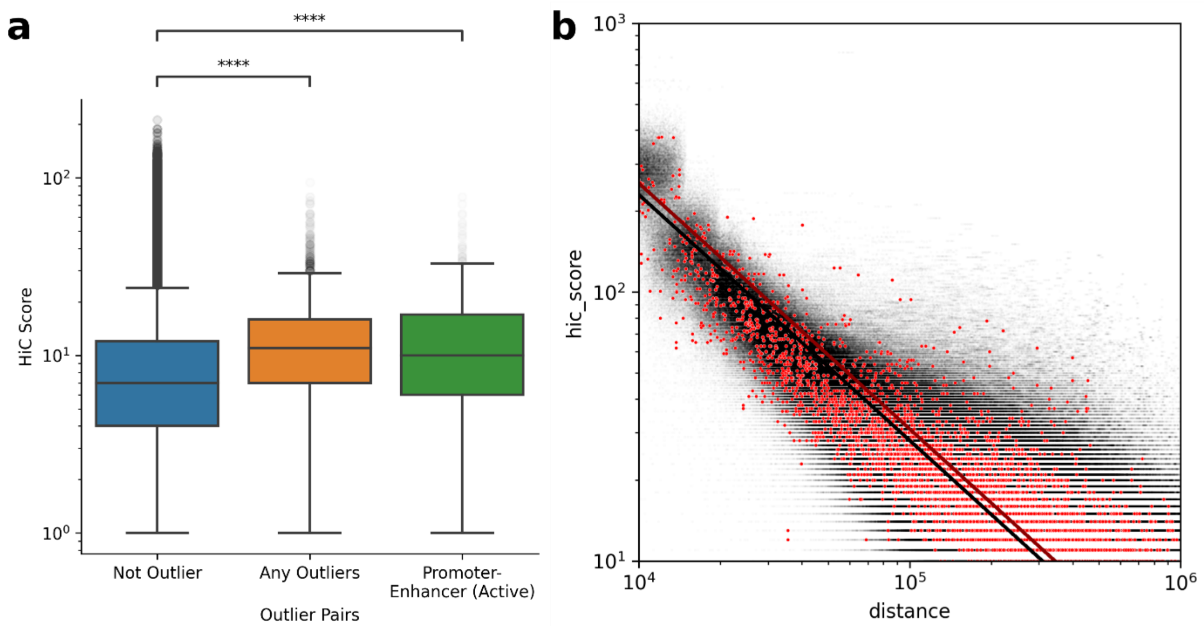


**Extended Data Fig. 5: Interaction between outlier pairs. (a)** Hi-C score distribution of accessible regions, which are at least 100 kilo-bp apart, by outlier status and annotation. P-values were calculated with the U-Mann Wily test and corrected for multiple testing using the Bonferroni method. **(b)** Log-log shows the Hi-C scores of non-outlier pairs (in black) and outlier pairs (in red). Hi-C scores decay by power law with increasing distances. We fit a power regression (indicated by black and red lines) on the data to test the interaction between outlier pairs while using distance as a control variable.

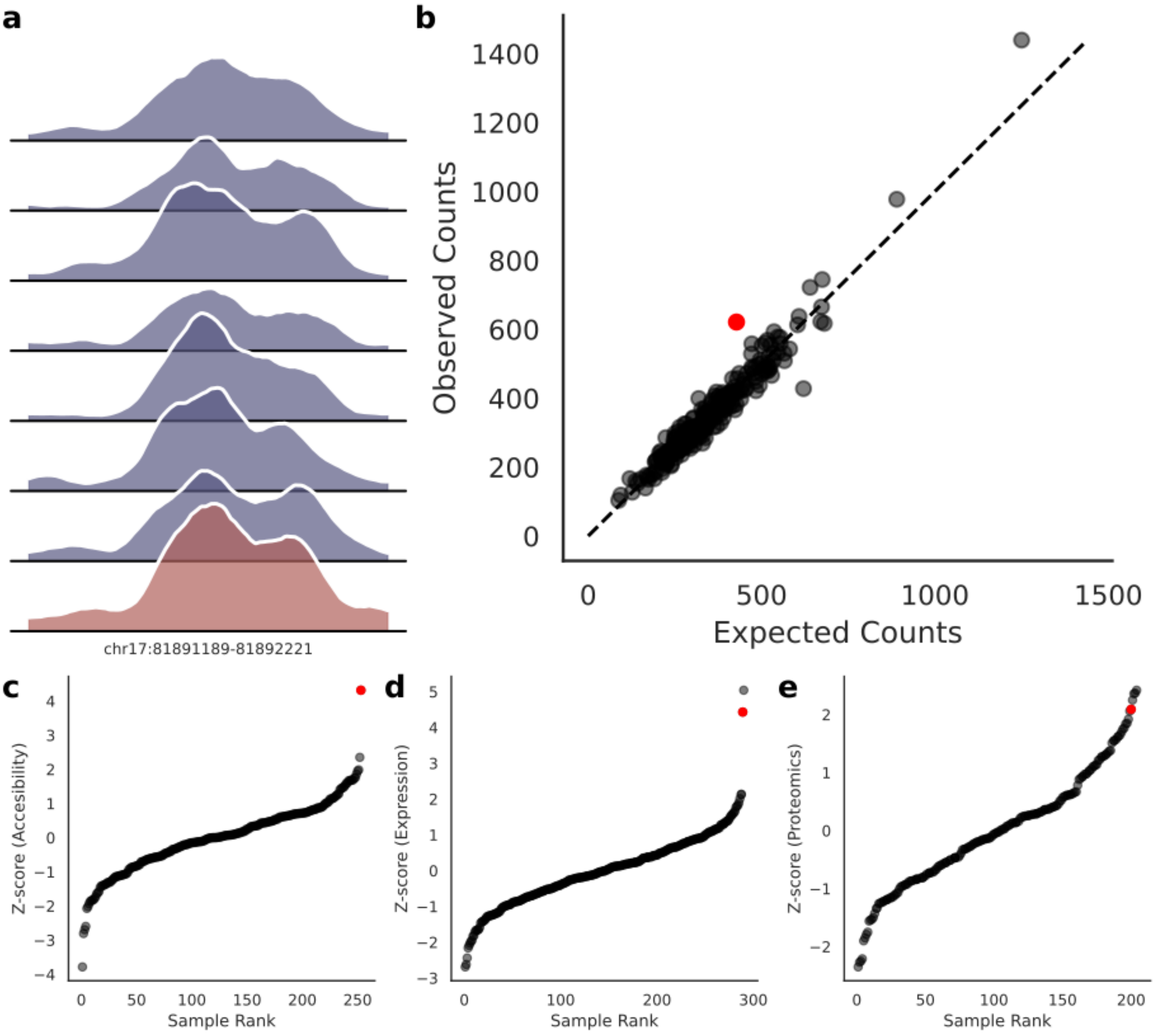


**Extended Data Fig. 6: Aberrations observed at multiple omics levels of *ALYREF* gene (a)** ATAC-seq read coverage at the promoter of the gene. The sample with an outlier promoter is indicated by red across panels. **(b)** Expected and observed accessibility in the promoter of the gene across samples **(c)** Z-score distribution of promoter accessibility **(d)** gene expression **(e)** protein levels across samples.

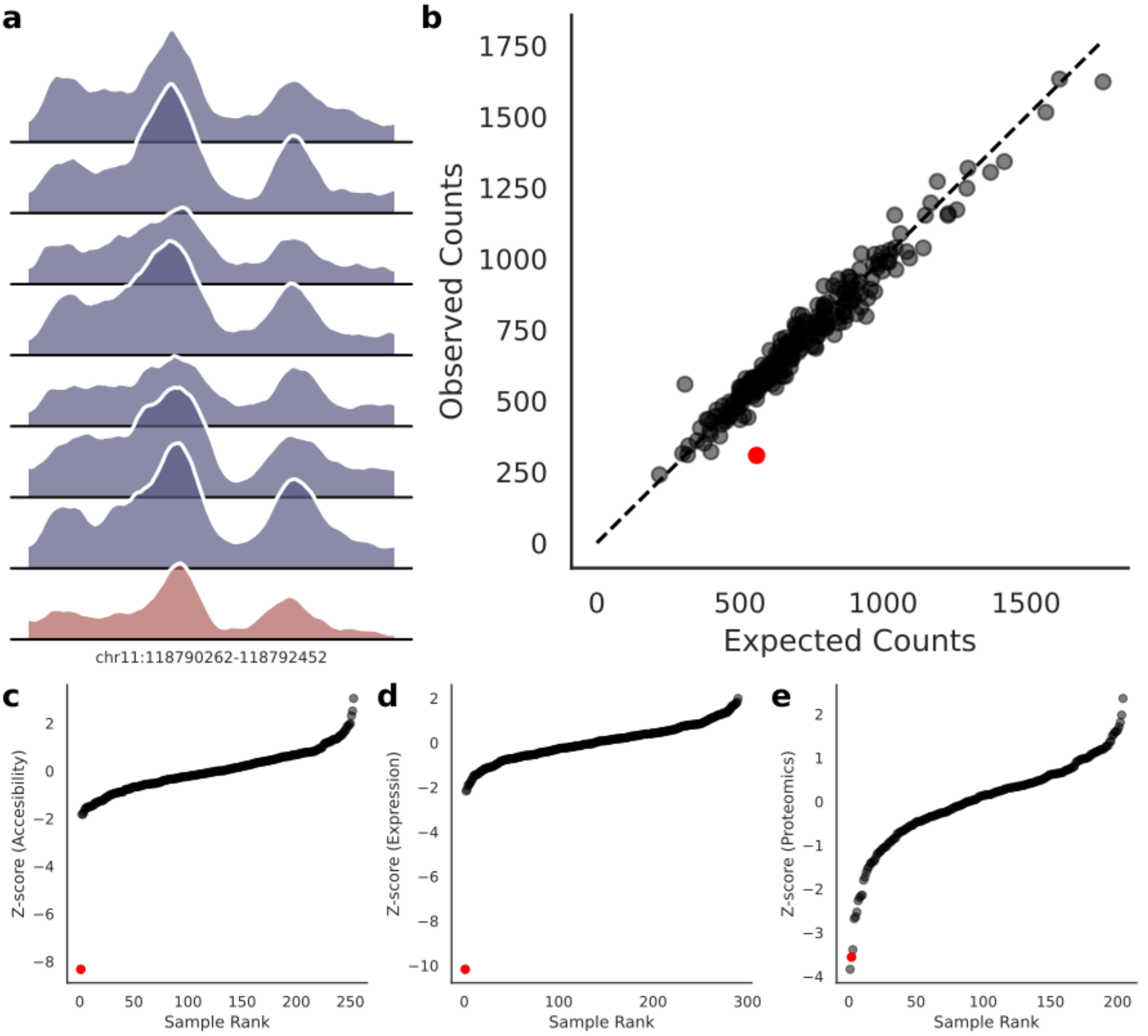


**Extended Data Fig. 7: Aberrations observed at multiple omics levels of *DDX6* gene (a)** ATAC-seq read coverage at the promoter of the gene. The sample with an outlier promoter is indicated by red across panels. **(b)** Expected and observed accessibility in the promoter of the gene across samples **(c)** Z-score distribution of promoter accessibility **(d)** gene expression **(e)** protein levels across samples.

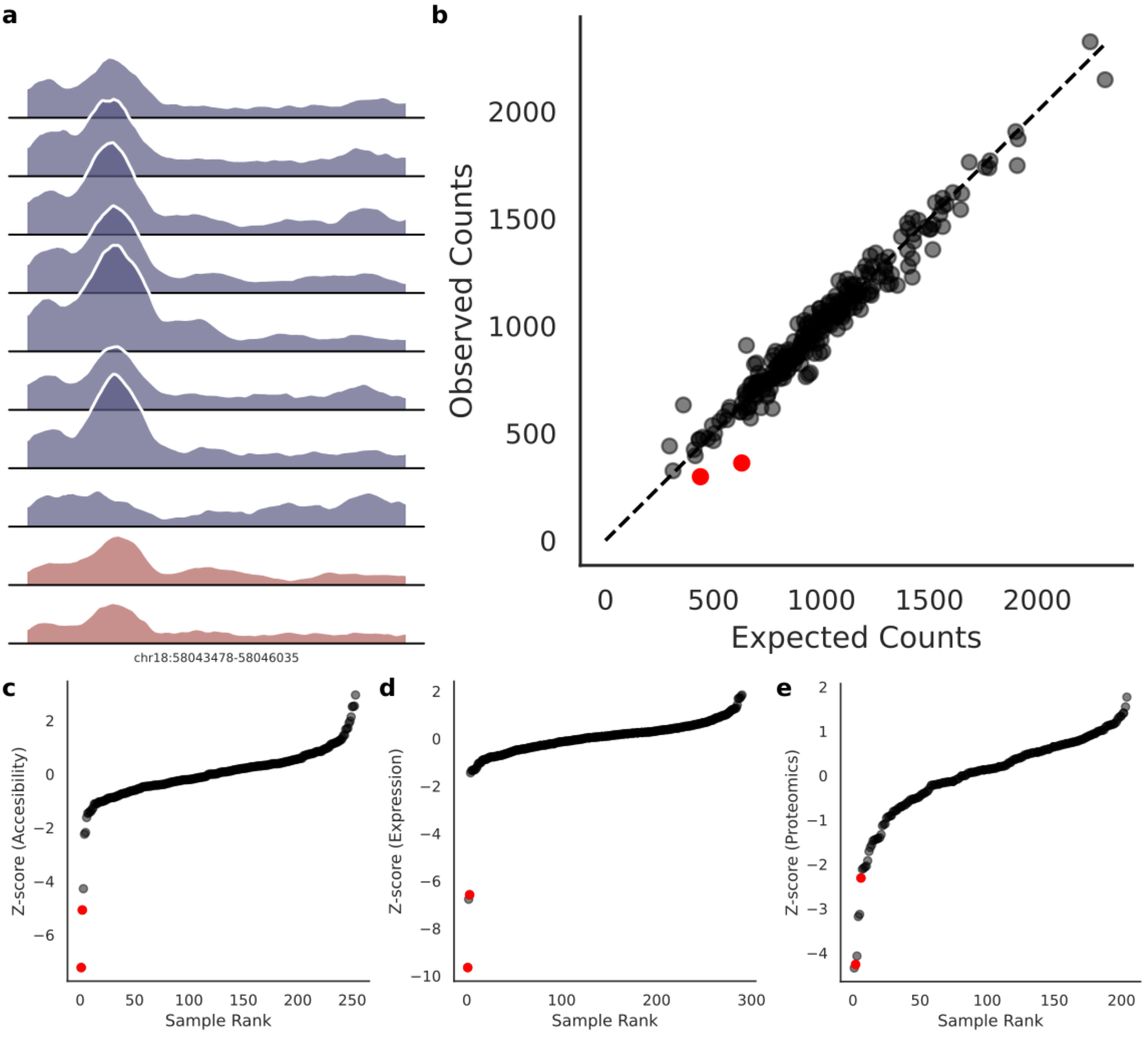


**Extended Data Fig. 8: Aberrations observed at multiple omics levels of *NEDD4L* gene (a)** ATAC-seq read coverage at the promoter of the gene. Samples with an outlier promoter are indicated by red across panels. **(b)** Expected and observed accessibility in the promoter of the gene across samples **(c)** Z-score distribution of promoter accessibility **(d)** gene expression **(e)** protein levels across samples.

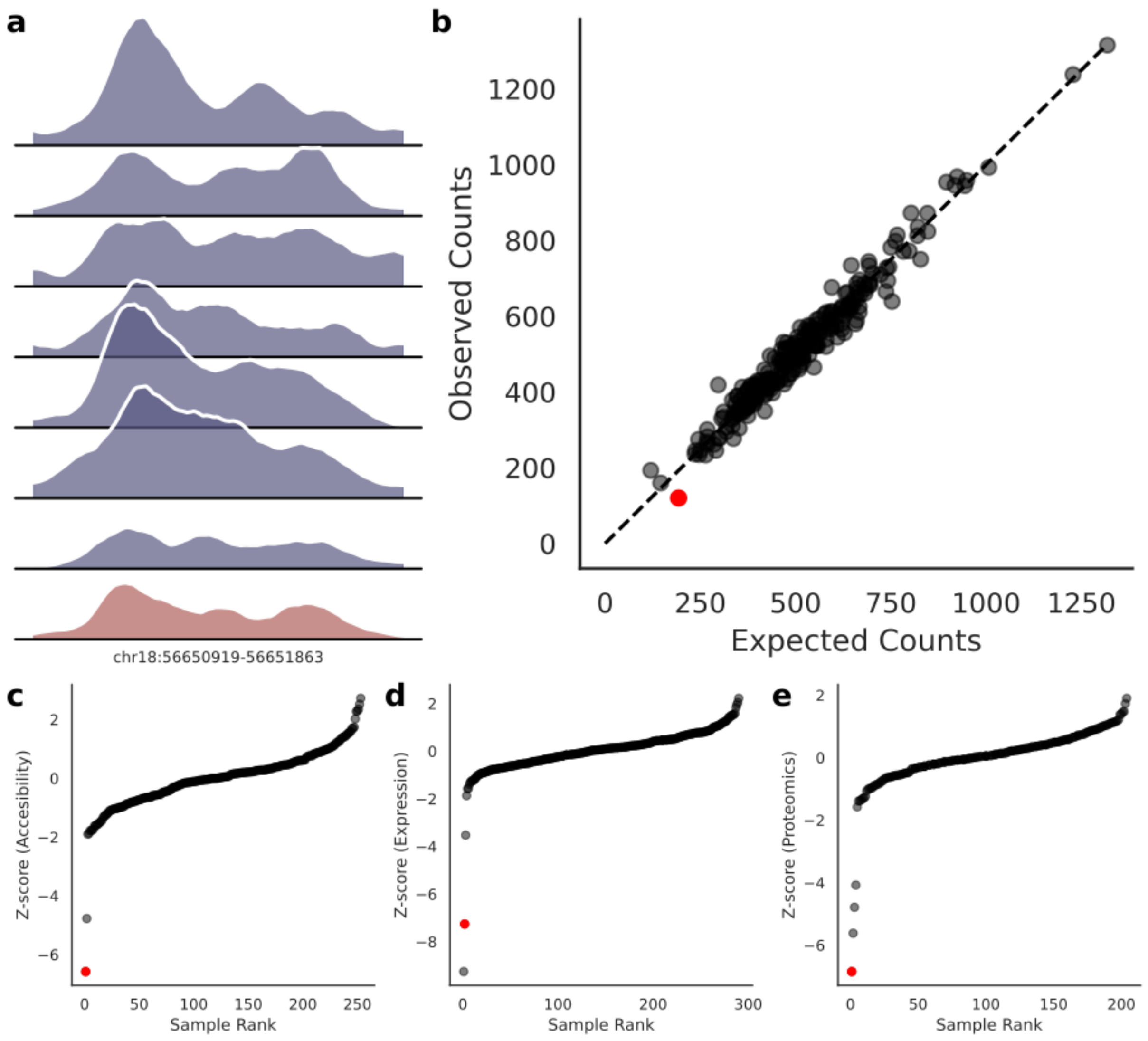


**Extended Data Fig. 9: Aberrations observed at multiple omics levels of *TXNL1* gene (a)** ATAC-seq read coverage at the promoter of the gene. The sample with an outlier promoter is indicated by red across panels. **(b)** Expected and observed accessibility in the promoter of the gene across samples **(c)** Z-score distribution of promoter accessibility **(d)** gene expression **(e)** protein levels across samples.

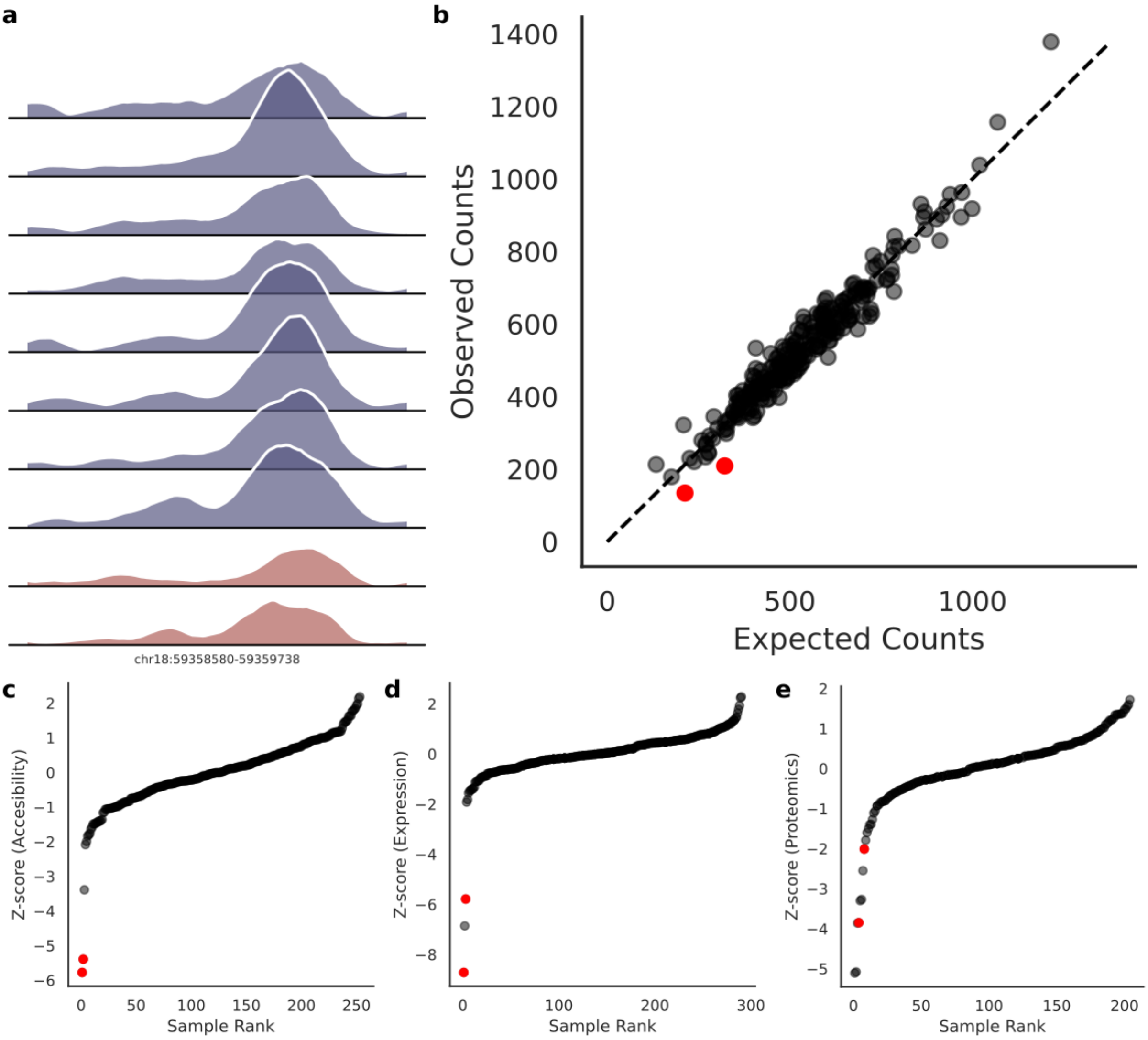


**Extended Data Fig. 10: Aberrations observed at multiple omics levels of *LMAN1* gene (a)** ATAC-seq read coverage at the promoter of the gene. Samples with an outlier promoter are indicated by red across panels. **(b)** Expected and observed accessibility in the promoter of the gene across samples **(c)** Z-score distribution of promoter accessibility **(d)** gene expression **(e)** protein levels across samples.

### Supplementary Algorithm

---

**Algorithm 1** Read counting algorithm for accessible regions

---

**Require:** a bam file and non-overlapping *peaks* sorted by position
**Ensure:** Read *counts* as a dictionary

**for each:** $chrom \in chromosomes$
  $peaks \leftarrow$ create a stack by subsetting peaks in $chrom$
  $reads \leftarrow$ create a stack by fetching reads from bam file for $chrom$

  $read \leftarrow reads.dequeue()$
  $peak \leftarrow peaks.dequeue()$

  **do**
    **if** $read.end <= peak.start$ **then**
      $read \leftarrow reads.dequeue()$
    **else if** $read.start >= peak.end$ **then**
      $peak \leftarrow peaks.dequeue()$
      $counts[peak] \leftarrow 0$
    **else**
      $counts[peak] + +$
      $read \leftarrow reads.dequeue()$
    **end if**
  **while** peaks $\neq \emptyset$

---
